# Stage-Specific Regulation of DNA Damage Repair by the Circadian Regulator, CRY1, in Prostate Cancer

**DOI:** 10.64898/2026.05.19.726303

**Authors:** Arwa Fallatah, Stefan DiFazio, Lakshmi Ravindranath, Thomas Janas, Filippo Pederzoli, Laila Scroggins, Noor Eldeen Abdalla, Orly Richter, Sally Elsamanoudi, Amina Ali, Jiji Jiang, Isabell Sesterhenn, Cara Schafer, Massimo Loda, Gregory Chesnut, Xiaofeng A. Su, Christopher M. McNair, Karen E. Knudsen, Ayesha A. Shafi

## Abstract

Circadian dysregulation is increasingly linked to prostate cancer (PCa) progression, yet its role in directing DNA damage response (DDR) pathway selection remains poorly understood. Here, we identify circadian cryptochrome 1 (CRY1), a core circadian regulator, as a stage-specific determinant of DDR dependencies. Integrated transcriptomic and CRISPR-based analyses reveal that CRY1 promotes non-homologous end joining (NHEJ) and base excision repair (BER)-associated programs in hormone-sensitive disease (HTS), while driving a switch toward homologous recombination (HR) dependency in castration-resistant prostate cancer (CRPC). Mechanistically, CRY1 couples proliferative signaling to genome maintenance, enabling tumor cells to tolerate genotoxic stress and sustain progression. Notably, loss of CRY1 exposes distinct, context-dependent DDR vulnerabilities, revealing repair plasticity as a targetable actionable feature of disease evolution. These findings position CRY1 as a central regulator of DDR rewiring and support CRY1-directed combination strategies with DDR inhibitors as a rationale to delay or prevent progression to advanced, treatment-resistant PCa.

## Introduction

Prostate cancer (PCa) is among the most commonly diagnosed malignancies in men and remains a major cause of cancer-related mortality^1^. While locally advanced is often responsive to androgen deprivation therapy (ADT), progression is frequent, and many tumors eventually evolve into castration-resistant prostate cancer (CRPC), an advanced state characterized by poor prognosis and limited therapeutic durability^2,3^. Despite the availability of next-generation androgen receptor (AR)-targeted therapies and cytotoxic agents, disease progression and treatment resistance remain pervasive clinical challenges. A fundamental gap in the field is the limited understanding of the molecular mechanisms that drive PCa progression across disease stages, particularly those that enable tumor cells to withstand therapeutic stress.

An emerging biological axis implicated in cancer development and progression is the disruption of the circadian clock^4,5^. The circadian system is an evolutionarily conserved molecular network that synchronizes cellular processes with environmental light-dark cycles, thereby coordinating proliferation, cell-cycle progression, and stress responses^5,6^. Epidemiological and experimental studies increasingly link circadian disruption to PCa risk and adverse clinical outcomes^4,5,7^. Beyond its role in maintaining temporal homeostasis, the circadian clock interfaces directly with pathways that preserve genome integrity, positioning circadian dysregulation as a potential driver of genomic instability and tumor evolution^8^.

Cryptochrome 1 (CRY1) is a core component of the circadian clock that functions as a transcriptional repressor within the molecular feedback loop^5,7^. Accumulating evidence indicates that CRY1 also exerts non-canonical functions in cancer, where it displays context-dependent oncogenic activity^7^. In PCa, elevated CRY1 expression has been associated with aggressive disease and poor clinical outcomes^7^. More recently, CRY1 has been identified as a modulator of the DNA damage response (DDR), a signaling network essential for detecting and repairing DNA lesions induced by endogenous stress and anticancer therapies^7^. Molecular and pharmacological suppression of CRY1 impairs PCa cell growth, induces G2/M cell-cycle arrest, and disrupts homologous recombination (HR) mediated DNA repair, suggesting that CRY1 contributes to tumor cell survival by maintaining DDR competence^7^.

Despite these insights, how CRY1 shapes DNA repair pathway choice during PCa progression remains incompletely understood. In particular, it is unknown whether CRY1-dependent regulation of the DDR is conserved or altered as tumors transition from hormone therapy sensitive (HTS) disease to castration-resistant states. Given that therapeutic resistance in CRPC is frequently accompanied by heightened replication stress and reliance on specific DNA repair mechanisms, defining the stage-specific functions of CRY1 within DDR networks is of critical importance.

In this study, we systematically investigated the role of CRY1 in regulating DNA repair pathway dependencies across PCa disease stages. Using genome-wide CRISPR screening approaches in HTS and CRPC PCa models, we identified DDR factors and repair pathways functionally modulated by CRY1 in distinct disease contexts. Our findings uncover a previously unappreciated role for CRY1 in directing DNA repair pathway choice during PCa progression and provide mechanistic insight into how circadian clock disruption contributes to therapeutic vulnerability in advanced disease.

## Results

### CRY1 is Pro-tumorigenic and Inhibition Decreases Growth in PCa

Previous studies have shown that gene amplification is frequently associated with increased gene expression^9^, and that oncogene amplification can emerge during cancer progression in a context-dependent manner with variable functional consequences across tumor types^10^. To assess the relevance of CRY1 dysregulation during disease evolution, publicly available PCa datasets were interrogated through cBioPortal, encompassing primary and metastatic prostate adenocarcinoma cohorts across localized disease, hormone therapy sensitive (HTS), castration-resistant prostate cancer (mCRPC), and neuroendocrine prostate cancer (NEPC)^11-14^ (**Fig. 1A**). CRY1 alteration frequency, including amplifications and mutations, increased alongside advanced disease stage consistent with selective pressure favoring CRY1 dysregulation during tumor progression. In parallel, enrichment of CRY1 mutational events in advanced cohorts (**Fig. 1A**) was consistent with the higher mutational burden that was observed in those metastatic cases compared to primary tumors and was indicative of aggressive clinical biology and poor prognosis^10^.

**Figure 1.**
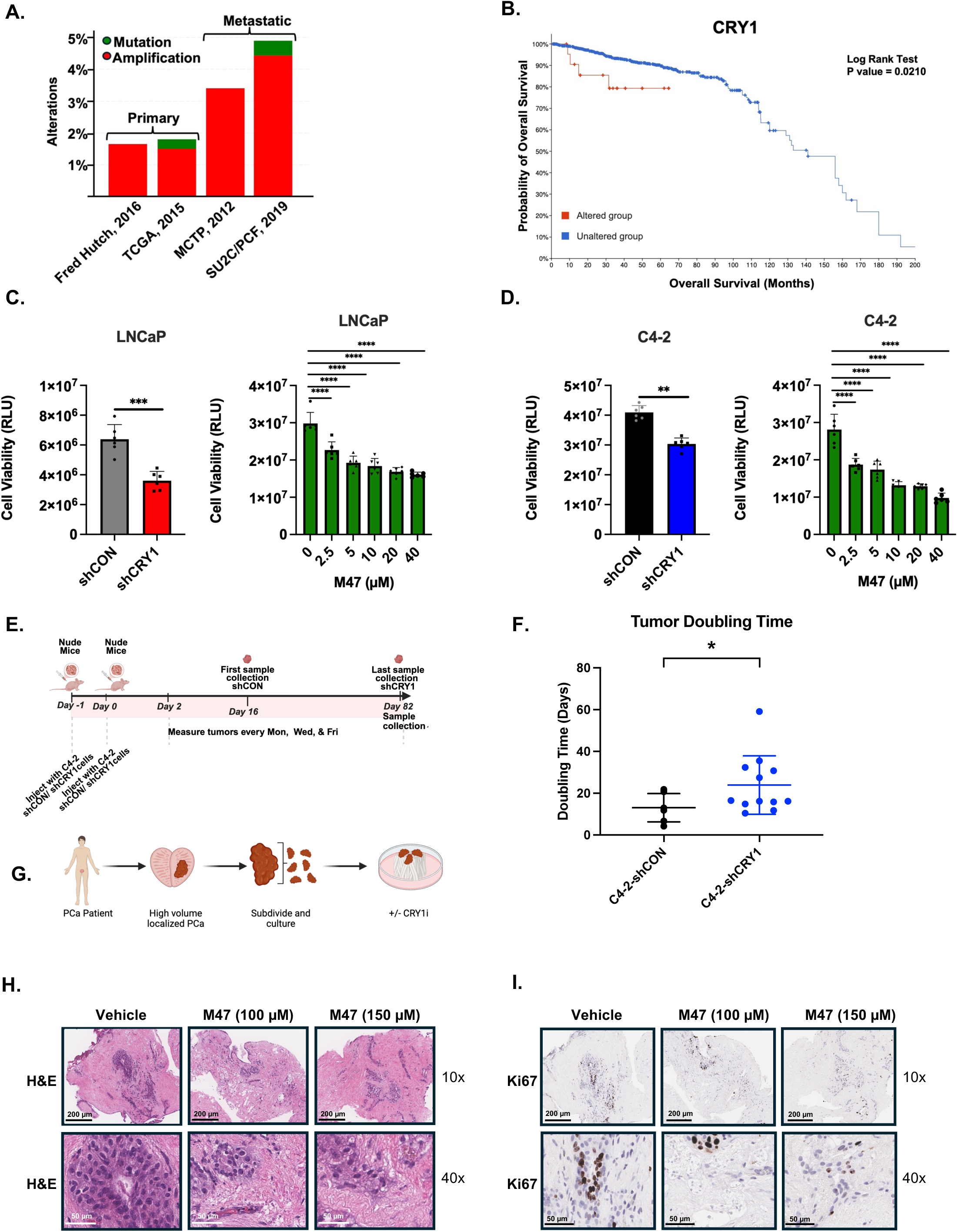
CRY1 is Protumorigenic and Inhibition Decreases Growth in PCa. **A.** Frequency of CRY1 gene alterations (i.e., amplifications and mutations) in primary and metastatic PCa datasets from cBioPortal—Fred Hutchinson CRPC (Nat Med 2016), TCGA(Cell 2015), MCTP(Nature 2012), and SU2C/PCF Dream Team (PNAS 2019) datasets. **B.** cBioPortal analyses for overall survival in PCa patients with CRY1 altered versus unaltered groups. **C.** Cell Titer-Glo 2.0 Cell Viability Assay in LNCaP shCRY1 and upon treatment with CRY1 inhibitor (M47). n=3, ***p<0.001 and ****p<0.0001. **D.** Cell Titer-Glo 2.0 Cell Viability Assay in C4-2 shCRY1 and upon treatment with CRY1 inhibitor (M47). n=3, **p<0.01 and ****p<0.0001. **E.** Schematic diagram of experimental design for C4-2 shCRY1 xenograft model. **F.** Tumor doubling time is significantly increased in C4-2-shCRY1 xenograft model Confirming that CRY1 promotes tumor growth in xenograft model, p<0.05. **G.** Patient-Derived Explants (PDE) prostate cancer tumor tissue treated with M47 (0, 100, and 150 uM) for 72 h and analyzed for **H-I.** H&E & Ki67 staining within PDE tissues. Scale bars are 50 & 200 µM.

To further evaluate the clinical relevance of CRY1 dysregulation, aggregate analysis across 1,987 samples derived from multiple independent studies pooled from different cohorts revealed that altered CRY1 expression was significantly associated with reduced overall survival (p=0.021; **Fig. 1B**), supporting an association between CRY1 dysregulation and adverse clinical outcome. Given this association, further studies examining whether CRY1 is functionally required for tumor cell survival across disease stages were performed using both HTS and CRPC models. Previous studies from this group demonstrated that CRY1 depletion inhibits proliferation within CRPC models¹, with this effect further amplified following irradiation. To assess the role of CRY1 at earlier disease stages, the LNCaP cell line was utilized to evaluate the effect of CRY1 on the androgen-dependent stage of the disease. We found that genetic depletion of CRY1 or pharmacological inhibition using the CRY1 inhibitor, M47, significantly reduced cell viability^15^ (**Fig. 1C**), indicating that CRY1 is required for survival in AR-driven PCa. This dependency persisted in CRPC models (i.e., C4-2 cells), where both CRY1 knockdown via shRNA and inhibition via M47 treatment markedly decreased viability (**Fig. 1D**), consistent with our previous observations in CRPC models via genetic perturbations^7^. Notably, CRPC cells exhibited increased sensitivity to CRY1 inhibition suggesting an increased dependency on CRY1 with disease progression. Importantly, CRY1 inhibition with M47 treatment did not exhibit any inhibitory effect on the normal prostate epithelial cell line, RWPE-1, indicating that its activity is selective for PCa cells while sparing non-malignant cells, highlighting CRY1 as a potential therapeutic target (**Supplementary Fig. 1A**). Collectively, these findings demonstrate that CRY1 dependence persists following the transition to androgen independence, but the underlying mechanism remains to be elucidated.

To confirm the oncogenic role of CRY1 on cellular growth is reincorporated in other experimental models, *in vivo* and *ex vivo* xenografts along with patient-derived explant tissue models were utilized, respectively. Specifically, to discern if CRY1 loss affects tumor growth *in vivo*, we established xenografts using C4-2 cells with CRY1 knockdown using previously reported doxycycline-inducible cell lines including C4-2-shCON and C4-2-shCRY1^7^. Briefly, doxycycline water (2mg/mL, 5% sucrose) was administered after tumor establishment. CRY1 depletion significantly delayed tumor growth compared with control (**Fig. 1E**), as reflected by a marked increase in tumor doubling time (**Fig. 1F**). Consequently, CRY1-depleted tumors required substantially longer to reach endpoint criteria defined as tumor size exceeding 1000 mm^3^, despite the time required for tumor development was not significantly different between groups (**Supplementary Fig. 1B**), demonstrating that CRY1 is required for efficient tumor expansion *in vivo*. These *in vivo* findings further support a critical role for CRY1 in maintaining tumor growth under physiological conditions, where proliferative demands must be balanced with ongoing genotoxic stress.

Translational relevance of CRY1 inhibition in human tumors was further assessed using patient-derived explants (PDE) from human PCa tissue following radical prostatectomy treated *ex vivo* with CRY1 inhibitor, M47, as described previously^15,16^. A minimum of three independent biological tissue samples were analyzed per condition, and representative results are shown. Briefly, PDEs were treated with 100 µM and 150 µM of M47 for 72 hours and refreshed daily for the duration of the treatment. CRY1 inhibition resulted in a dose-dependent reduction in proliferation, as assessed by Ki67 staining (**Fig. 1G-I**). A previously published study has shown that the CRY1 inhibitor, M47, has a favorable toxicity profile *in vivo*, as it did not affect mouse body weight *in vivo* and is capable of crossing the blood brain barrier^15^, supporting continued therapeutic investigation and its potential translational application in PCa patients. Collectively, these findings identify CRY1 as a clinically relevant regulator in PCa aggressiveness and demonstrate a functional dependency that persists across disease states.

### CRY1 Loss Induces Stage-specific Transcriptional Rewiring of DNA Damage Repair Related Pathways

Given prior evidence that CRY1 promotes PCa growth and associates with metastatic disease exhibiting more alteration events, we hypothesized that CRY1 executes stage-specific functions as the tumors progress. To test this, we performed a genome-wide analysis of CRY1-sensitive transcriptional networks using newly generated, doxycycline-regulated isogenic pairs with inducible CRY1 knockdown in LNCaP (HTS) cells and compared the resulting transcriptomes with C4-2 (CRPC) profiles reported previously^7^. Cells were treated with doxycycline for 72 hours prior to RNA sequencing. Efficient CRY1 knockdown was confirmed by examining levels of protein via Western blot analysis, with approximately 52% reduction of CRY1 protein following shRNA induction (**Fig. 2A**, left), and strong concordance among biological replicates (**Supplementary Fig. 2A**).

**Figure 2.**
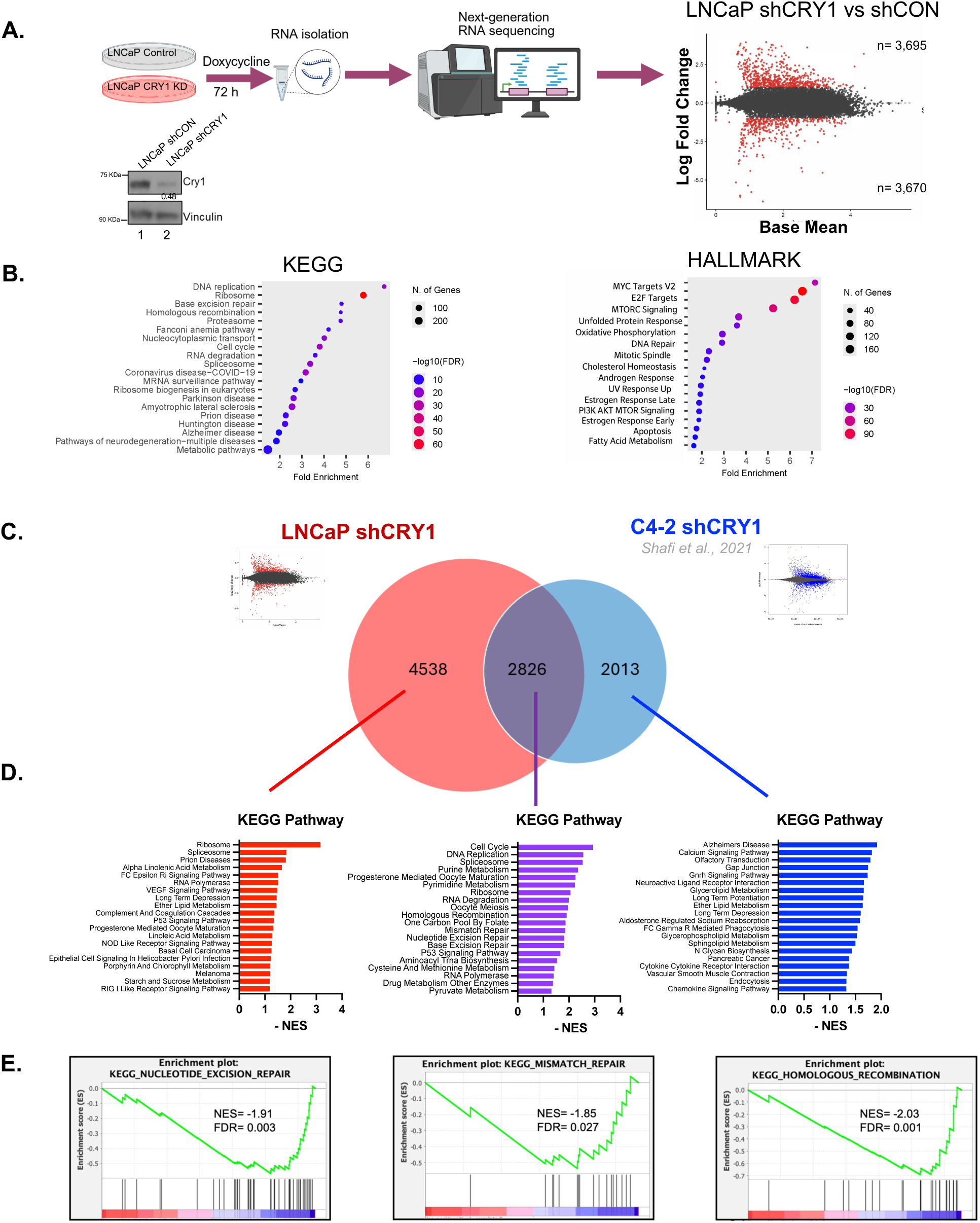
Altered CRY1 Expression Correlates with Aggressive Disease. **A-B.** Workflow showing RNA-seq analysis after CRY1 knockdown in LNCaP cells, MA plot displaying differential gene expression, and Western blot confirming CRY1 knockdown. (A) Pathway enrichment analysis differentially expressed genes in LNCaP cells, KEGG and Hallmark. (B) **C.** Venn diagram comparing differentially expressed genes between LNCaP and C4-2 cells after CRY1 knockdown, and KEGG pathway enrichment for unique and shared gene sets using Enrichr. p<0.01 **D-E.** Three Gene Set Enrichment Analysis (GSEA) KEGG pathways and leading-edge plots highlighting specific DDR-related pathways (e.g., base excision repair, mismatch repair, and homologous recombination) that are significantly enriched after CRY1 knockdown.

CRY1 loss triggered broad transcriptional reprogramming, with 3,695 genes upregulated and 3,670 genes downregulated at an adjusted p< 0.05, indicating that CRY1 influences large, coordinated gene networks (**Fig. 2A**, right). Gene set enrichment analysis (GSEA) using ShinyGo revealed robust enrichment of pathways central to cancer biology, including DNA replication, oxidative phosphorylation, metabolism, hormonal signaling, and multiple DNA-repair processes (**Fig. 2B**), consistent with a model in which CRY1 orchestrates core replication/repair circuitry in a context-dependent manner across HTS and CRPC states^7^. Loss of CRY1 reprogrammed the transcriptome in a stage-selective manner. In LNCaP, 4,538 genes were uniquely deregulated, whereas 2,013 genes were uniquely affected in C4-2 cells. Importantly, 2,826 transcripts were shared between models, which highlights the core biological pathways regulated by CRY1 independently from the disease stages (**Fig. 2C**). This partial conservation with progressive rewiring is consistent with known molecular evolution from hormone-treatment sensitive to CRPC^17^.

To further dissect the biological programs underlying these shifts, we performed GSEA using genes uniquely differentially expressed in LNCaP or C4-2 cells, as well as genes commonly altered between both models (**Fig. 2D**). KEGG and Hallmark GSEA revealed that CRY1 knockdown in early-stage PCa, modeled by LNCaP cells, induces broad transcriptional reprogramming across ribosome, spliceosome, metabolic, and immune-related pathways, consistent with a role for CRY1 in maintaining cellular homeostasis (**Fig. 2D and Supplementary Fig. 2B**). Future studies investigating the role of CRY1 in rewiring the metabolism in PCa as the disease progress are ongoing. As disease advances to the castration-resistant state, modeled by C4-2 cells, CRY1 knockdown reveals both conserved activation of cell cycle, DNA replication, spliceosome, and core metabolic pathways and CRPC-specific enrichment of inflammatory and cytokine-mediated signaling, collectively highlighting dysregulation of DDR associated pathways as a central feature of CRY1-dependent regulation across PCa progression (**Fig. 2D and Supplementary Fig. 2C-D**). Notably, pathways commonly altered upon CRY1 knockdown in both LNCaP and C4-2 cells are enriched for cell cycle regulation, DNA replication, spliceosome function, and nucleotide and energy metabolism (Fig. 2D and Supplementary Fig. 2C), underscoring a conserved reliance on core DDR-linked processes that persist across the disease continuum, highlighting a conserved reliance on genome maintenance pathways across PCa progression^7,18^.

GSEA assessment revealed that the base excision repair (BER) pathway was significantly upregulated within the LNCaP-unique gene set (FDR <0.003), indicating robust activation of BER programs in the HTS state. In contrast, HR was specifically enriched in C4-2 cells (FDR <0.001), reflecting the repair landscape of CRPC. Notably, mismatch repair (MMR) emerged as a significantly upregulated pathway within the shared transitional gene set between HTS and CRPC (FDR < 0.027), suggesting that MMR represents an intermediate repair program engaged during the HTS-to-CRPC transition, a phenomenon consistent with emerging models of adaptive DDR pathway switching during cancer evolution^19-21^.

The role of HR in CRPC has been addressed previously, where CRY1 was shown to mediate DNA-repair programs during disease progression and to associate with poor clinical outcome, nominating CRY1 as an emerging target for investigation^7^. Building on this foundation, the present data indicating that CRY1-dependent circuitry is stage-contingent where in HTS PCa, CRY1 aligns with BER-linked transcriptional and metabolic modules. Conversely, in CRPC, CRY1 increasingly interfaces with HR-directed repair and growth pathways. This state-specific alignment suggests that CRY1 functions not merely as a general stress adaptor but as a contextual organizer of genome-stability and other cancer driving programs.

Together, these observations support a model in which CRY1 acts as a stage-contingent transcriptional hub, steering distinct repair as BER in HTS transitioning to HR in CRPC as tumors evolve, and thereby shaping the molecular trajectory of PCa progression.

### CRY1 Loss Uncovers Stage-Specific DDR Dependencies in PCa

Given the stage-contingent transcriptional reprogramming of DDR pathways following CRY1 loss (**Fig. 2**), we hypothesized that CRY1 depletion functionally rewires genetic dependencies within DDR machinery across PCa disease states. To test this, we performed a focused, pooled CRISPR-Cas9 screen targeting core DDR genes in PCa models with inducible CRY1 knockdown compared to control knockdown (i.e., shCON). This approach leverages targeted CRISPR screening strategies previously shown to effectively resolve pathway-level DNA repair dependencies^22^ and are informed by established DDR-focused CRISPR library designs^23^. A pooled DDR CRISPR library targeting key DNA repair genes was introduced into LNCaP (HTS) and C4-2 (CRPC) cell lines harboring doxycycline-inducible shCRY1 or shCON constructs (**Fig. 3A**). This library design follows prior focused DDR CRISPR screening efforts that systematically target known and candidate DNA repair regulators to reduce complexity relative to genome-wide screens^23^. Following Cas9-mediated editing, cells were maintained under proliferative conditions, and sgRNA representation was quantified over time to identify genes whose disruption impaired cellular fitness upon CRY1 depletion. Targeted CRISPR approaches have proven effective for interrogating functional DNA repair dependencies while reducing the complexity and noise inherent to genome-wide screens^22^. Negative selection hits are defined as genes whose sgRNAs were significantly depleted over time (from passage zero (P0) to passage ten (P10)) and were identified using robust rank aggregation (RRA) analysis. Briefly, this was implemented using the MAGeCKFlute analysis framework, which integrates MAGeCK RRA-based gene-level scoring with normalization and quality-control procedures optimized for pooled CRISPR screens^24^.

**Figure 3.**
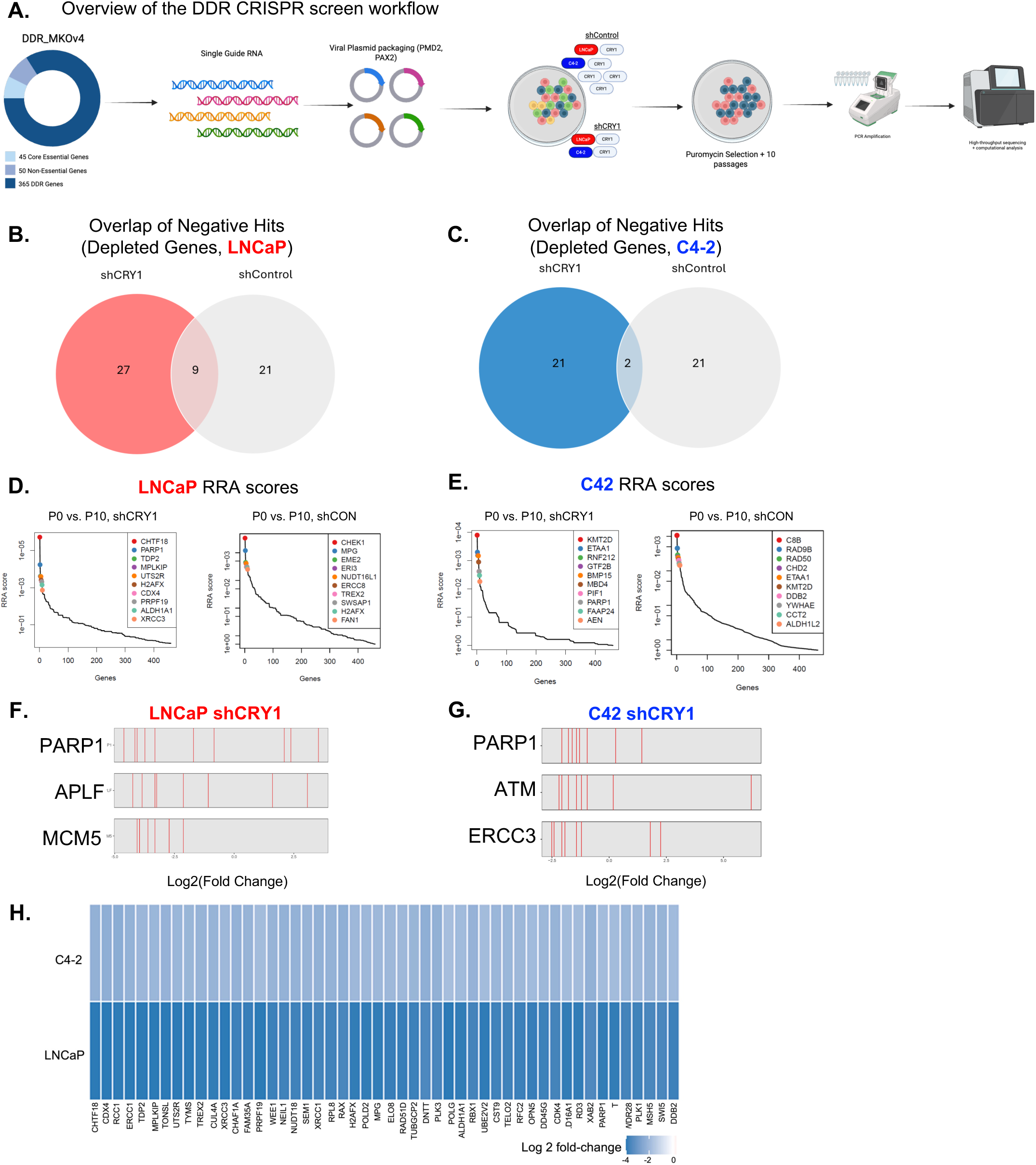
DNA Damage Repair CRISPR screen with CRY1 knock down. **A.** Library overview and schematic of DDR CRISPR screen workflow. **B.** Venn diagram showing overlap of negatively enriched hits (essential DDR genes) between CONtrol and CRY1 KD in LNCaP cells. **C.** Venn diagram showing overlap of negatively enriched hits (essential DDR genes) between CONtrol and CRY1 KD in C4-2 cells. **D.** Gene-level RRA enrichment plots highlighting top negatively selected DDR genes under CRY1 KD in LNCaP cells. **E.** Gene-level RRA enrichment plots highlighting top negatively selected DDR genes under CRY1 KD in C4-2 cells. **F-G.** Guide RNA plots for targets of interest in LNCaP (PARP1, APLF, MCM5) and in C4-2 (PARP1, ERCC3, ATM). **H.** Heatmap depicting Log2FoldChange for 20 commonly shared dependencies identified in LNCaP and C42, illustrating increased reliance on specific repair pathways following CRY1 KD.

Efficient and sustained CRY1 knockdown in both LNCaP and C4-2 cells across the screening time course was confirmed by Western Blot at P0 and P10, with appropriate loading controls (**Supplementary Fig. 3A-B**). In parallel, genomic DNA integrity and comparable library amplification between control and CRY1-depleted conditions were verified by agarose gel electrophoresis prior to sequencing (**Supplementary Fig. 3C-D**). Consistent sgRNA representation and reproducible read count distributions across biological replicates at both P0 and P10 further supported the technical robustness of the screen (**Supplementary Fig. 3E-L**). In LNCaP cells, CRY1 knockdown revealed a distinct pattern of genetic vulnerability within DDR pathways. Overlap analysis demonstrated both shared and CRY1-specific depleted genes, indicating that CRY1 loss unmasks additional essential DDR components in the hormone-sensitive disease state (**Fig. 3B**). RRA analysis further identified *PARP1*, *APLF*, and *MCM5* among the most significantly depleted hits selectively enriched upon CRY1 loss ( **Fig. 3D and F**), consistent with emerging evidence that discrete DNA repair nodes exhibit context-dependent essentiality^22^.

In contrast, CRISPR screening in C4-2 cells identified a partially overlapping but distinct CRY1-dependent dependency landscape. While a subset of DDR dependencies was conserved, *CRY1* depletion preferentially sensitized the CRPC model to loss of genes implicated in recombination-associated and replication-coupled DNA repair processes, including *PARP1*, *ATM*, and *ERCC3* (**Fig. 3C, E, and G**), suggesting increased reliance on HR pathways in advanced disease. These findings are consistent with prior functional studies demonstrating that loss of checkpoint and repair regulators can confer profound effects on cellular fitness and treatment response under conditions of genomic stress^24^.

Comparative analysis across models revealed a core subset of DDR genes depleted in both LNCaP and C4-2 cells, consistent with conserved DNA repair requirements across PCa states (**Fig. 3H**). However, the majority of CRY1-dependent negative selection hits were model-specific, underscoring the stage-selective nature of CRY1-organized repair circuitry. Such context dependency aligns with recent combinatorial CRISPR studies demonstrating that DDR genes operate within highly interconnected yet condition-specific functional networks^23^. Notably, the CRISPR-derived dependency patterns align closely with transcriptional programs defined in Figure 2. Genes linked to BER and replication stress responses were preferentially essential in HTS cells, whereas the CRPC model exhibited increased reliance on pathways associated with HR and complex DNA damage processing. Together, these results functionally validate the RNA-sequencing-derived model in which *CRY1* acts as a stage-contingent organizer of DNA repair pathway utilization. Collectively, these CRISPR screen data demonstrate that *CRY1* loss does not globally impair cell viability but instead reprograms dependency on discrete DNA repair modules in a disease-state-dependent manner. This functional stratification complements transcriptional profiling and reinforces a model in which CRY1 coordinates adaptive DDR pathway selection during PCa progression, nominating CRY1-associated repair nodes as candidate therapeutic vulnerabilities.

### CRY1-Dependent Gene Depletion Reveals Shared and Disease-State Specific DDR Pathways in PCa Models

To connect functional genetic dependencies with transcriptional consequences of CRY1 depletion, we integrated significantly depleted hits from the DDR-focused CRISPR screen performed in LNCaP and C4-2 cells. This analysis revealed a restricted but biologically coherent overlap in DDR dependencies between the two models. In total, 32 genes were uniquely depleted in LNCaP cells, 20 genes were uniquely depleted in C4-2 cells, and 3 genes (*PARP1, AEN*, and *KMT2D*) were depleted in both models (**Fig. 4A**), underscoring limited conservation of DDR vulnerabilities across disease-states^7,25-30^. In line with the limited overlap observed in the Venn analysis, Maximum Likelihood Estimation (MLE) plots highlighted distinct, disease-state specific DDR dependencies following CRY1 loss. In LNCaP cells, the strongest negative selections included *PARP1, APLF*, and *MCM5*, genes with established roles in nucleotide excision repair (NER), replication-associated repair, and genome maintenance, consistent with preferential engagement of NER-linked programs in early-stage PCa^31^ (**Supplementary Fig. 4A**). In contrast, C4-2 cells showed prominent depletion of *ATM, PARP1*, and *ERCC3*, aligning with the enrichment of HR-related pathways in castration-resistant disease^20,32-34^ (**Supplementary Fig. 4B**). ATM is a central regulator of double-strand break signaling and HR pathway fidelity, and its loss is prevalent in advanced PCa and associated with altered therapeutic sensitivity ^20,32-34^. *ERCC3*, a core TFIIH helicase, has been linked to transcription-associated DNA repair and genome stability in oncogenic stress states characteristic of CRPC^35,36^.

**Figure 4.**
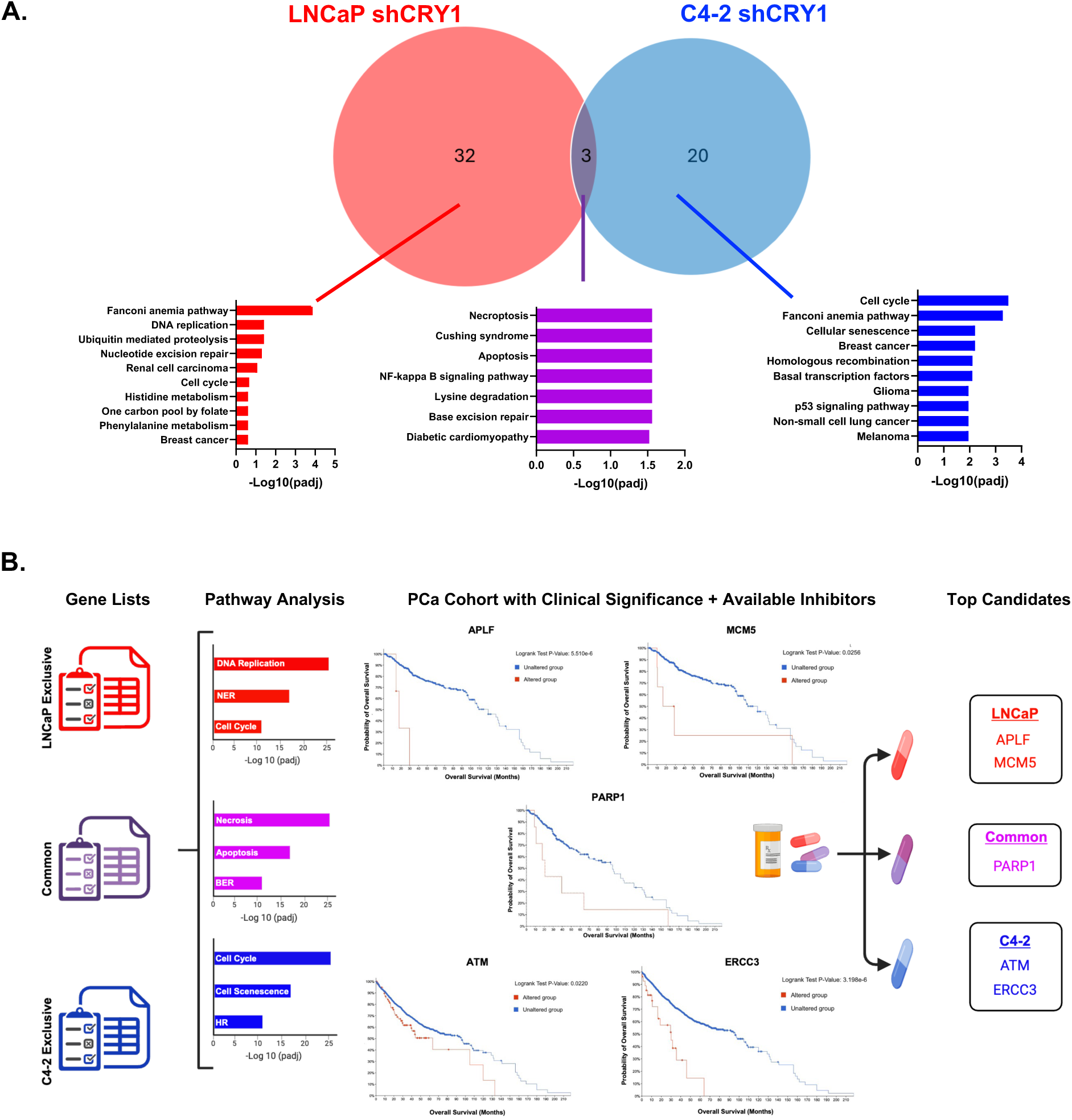
Comparative analysis of depleted gene overlap and DDR pathway analysis between LNCaP and C4-2 cell lines. **A.** Venn diagram illustrating overlap of significantly depleted genes (negative RRA selections, p <.05) between LNCaP and C4-2, with associated KEGG pathway enrichment. **B.** Schematic illustrating unique and shared pathways in HTS (LNCaP) and CRPC (C4-2) prostate cancer models, highlighting candidate DDR and metabolic vulnerabilities. The diagram depicts pathway nodes selectively altered in HTS (APLF and MCM5) and CRPC (ATM and ERCC3), along with available small-molecule inhibitors used or proposed for further functional validation. Kaplan–Meier analyses from cBioPortal are incorporated to compare overall survival in PCa patients with altered versus unaltered expression of APLF and MCM5 (HTS-specific targets) and ATM and ERCC3 (CRPC-specific targets), demonstrating the clinical relevance of these pathway perturbations.

To contextualize these dependencies, we performed KEGG pathway enrichment analysis using Enrichr. In agreement with transcriptomic analysis, DDR CRISPR screen results uncovered selective enrichment of NER-associated pathways in CRY1-depleted LNCaP cells, a shared enrichment of BER pathways across both models, and preferential enrichment of HR pathways in CRY1-depleted C4-2 cells (**Fig 4A**). Hallmark pathway analysis of negatively enriched CRISPR hits further validated the transcriptomic programs induced by CRY1 loss. LNCaP-specific dependencies were enriched for pathways linked to cellular homeostasis and proliferation (**Supplementary Fig 4C**), whereas genes commonly altered in both LNCaP and C4-2 cells converged on cell-cycle and core DDR programs (**Supplementary Fig 4D**). In contrast, C4-2-specific hits preferentially mapped to stress-responsive, apoptotic, and DNA repair associated pathways, consistent with the transcriptional signature of castration-resistant disease (**Supplementary Fig. 4E**). Individual genes lists are listed in (**Supplementary Fig. 4F**). Together, these findings integrate functional genetic dependencies with transcriptional remodeling to reinforce the Venn-based results, revealing a shared requirement for BER-associated repair alongside a progressive shift in DDR pathway engagement following CRY1 loss, wherein early-stage PCa cells preferentially deploy NER-linked repair programs, while castration-resistant cells transcriptionally and functionally rewire toward HR-dependent DNA repair during disease progression.

Next, the clinical relevance of CRY1-dependent DDR vulnerabilities was assessed by interrogating publicly available PCa cohorts through cBioPortal, integrating genetic alterations with patient survival data and the availability of corresponding pharmacologic inhibitors. This analysis refined candidate selection to a small, disease-state specific set of DDR genes. In the early-stage context represented by LNCaP cells, *APLF* and *MCM5* were associated with clinical outcome and prioritized as candidate dependencies. *PARP1* emerged as a shared vulnerability across both models, consistent with its central role in BER. In contrast, *ATM* and *ERCC3* were selectively associated with survival in the castration-resistant C4-2 model. Together, these analyses nominate *APLF* and *MCM5* for early-stage disease, *PARP1* as a common dependency, and *ATM* and *ERCC3* for CRPC as clinically relevant DDR targets downstream of CRY1 loss (**Fig. 4B**). Collectively, these findings consolidate functional genetic screening with transcriptional remodeling to reinforce Venn-based dependency analyses, revealing (i) a shared requirement for BER-associated repair and (ii) a progressive shift in DDR pathway engagement following CRY1 loss, wherein early-stage PCa cells preferentially deploy NER-linked repair programs, while castration-resistant cells transcriptionally and functionally rewire toward HR-dependent DNA repair during disease progression^7,25,32^.

### CRY1 Inhibition Differentially Sensitizes HTS and CRPC Models to DDR Inhibitors in a Context-Dependent Manner

To functionally validate stage-specific DDR dependencies identified by CRISPR screening, this study assessed sensitivity to pharmacologic inhibitors targeting shared and lineage-restricted DDR nodes. Notably, the concordant phenotypes observed across isogenic shCRY1 lines, CRY1-knockdown systems, and pharmacologic CRY1 inhibition demonstrate that the enhanced sensitization reflects on-target consequences of CRY1 loss, rather than off-target compound activity.

In LNCaP (HTS) isogenic lines, shCON cells showed the expected sensitivity to PARP inhibitors (olaparib, 10 µM, talazoparib, 5 µM; 72 hours), whereas shCRY1 cells exhibited significantly greater viability loss at matched doses (**Fig. 5A**). Treatment-response differences were evaluated using two-way ANOVA with multiple-comparison correction. Importantly, co-targeting of CRY1 and PARP resulted in greater suppression than either monotherapy (**Fig. 5B**). Treatment-response differences were evaluated using two-way ANOVA with multiple-comparison correction. A similar pattern emerged in C4-2 (CRPC) isogenic lines. In **Fig. 5C**, shCON cells responded as anticipated to PARP inhibition (olaparib, 1 µM, talazoparib, 1 nM; 72 hours), but shCRY1 cells showed a marked increase in inhibitor sensitivity. Treatment-response differences were evaluated using two-way ANOVA with multiple-comparison correction. In **Fig. 5D**, CRY1 knockdown produced maximal sensitization, and as observed in HTS cells dual CRY1 and PARP inhibition further enhanced viability loss in CRPC cells, underscoring CRY1’s central role in maintaining HR-dependent repair capacity across disease states. Treatment-response differences were evaluated using two-way ANOVA with multiple-comparison correction. These pharmacologic responses are consistent with CRY1’s emerging role as a regulator of DDR pathway choice and align with the canonical PARP1-driven BER and single-strand break (SSB) repair cascade, in which PARP1 detects DNA breaks and catalyzes PARylation to recruit XRCC1, POLβ, and LIG3 for efficient repair^26,27,29^. Notably, CRY1 inhibition has also been shown to sensitize ovarian cancer cells to PARP inhibition, reinforcing CRY1 as a context-dependent regulator of HR proficiency across tumor types^37^.

**Figure 5.**
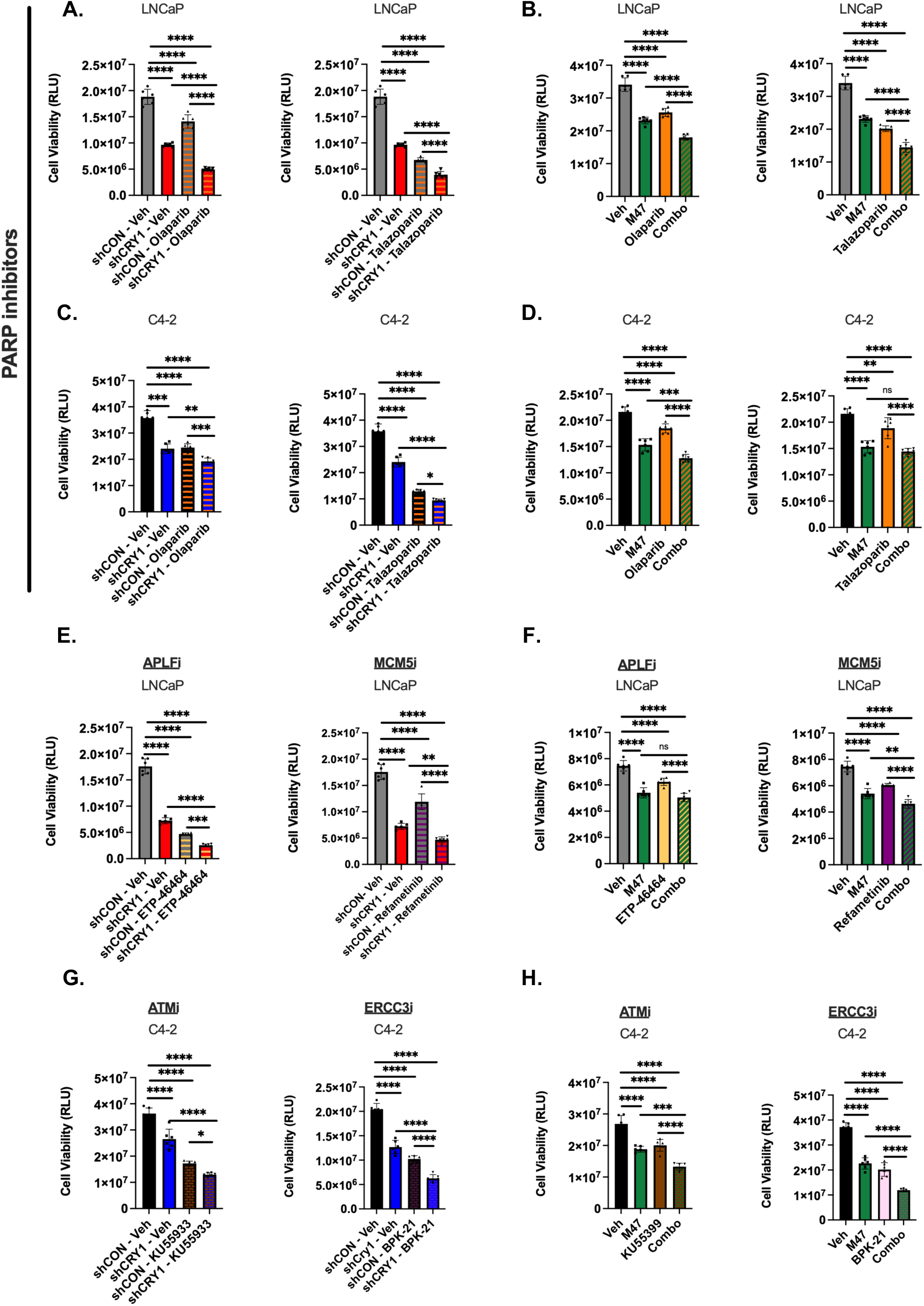
CRY1 Inhibition Differentially Sensitizes HTS & CRPC Models to Different DDR inhibitors in a Context Dependent Manner. **A.** Relative viability of LNCaP cell with CRY1 knockdown following treatment with PARP inhibitors, a class of DDR inhibitors common to both HTS and CRPC models, assessed by CellTiter-Glo assay. **B.** Relative viability of LNCaP cell treated with the CRY1 inhibitor M47 in combination with PARP inhibitors, measured by CellTiter-Glo assay. **C.** Relative viability of C4-2 cell with CRY1 knockdown following treatment with PARP inhibitors, a class of DDR inhibitors common to both HTS and CRPC models, assessed by CellTiter-Glo assay. **D.** Relative viability of C4-2 cell treated with the CRY1 inhibitor M47 in combination with PARP inhibitors, measured by CellTiter-Glo assay. **E.** Relative viability of LNCaP cells with CRY1 knockdown treated with HTS-specific DDR inhibitors targeting APLF and MCM5. **F.** Relative viability of LNCaP cells treated with M47 in combination with APLF and MCM5 inhibitors. **G.** Relative viability of C4-2 cells with CRY1 knockdown treated with CRPC-specific DDR inhibitors targeting ATM and ERCC3. **H.** Relative viability of C4-2 cells treated with M47 in combination with ATM and ERCC3 inhibitors. All viability measurements were obtained using the CellTiter-Glo luminescence assay. Bars show mean±SEM of three biological replicates. Statistical comparisons performed using one-way ANOVA. Asterisks indicate statistical significance between groups. *p<0.05, **p<0.01, ***p<0.001, and ****p<0.0001.

Disease-stage specific analyses identified *APLF*, a PBZ-domain chromatin anchor involved in non-homologous end-joining (NHEJ) and fork protection, and *MCM5*, a CMG-helicase component, as HTS-specific liabilities. Supporting this, ETP-46464, a potent mTOR and ATR inhibitor that also targets the APLF DNA-binding domain (0.1 µM, 72 h) were utilized to assess impact in PCa cell models (**Fig. 5E**) Similarly, refametinib (BAY 86-9766), a selective non-ATP-competitive allosteric inhibitor of MEK1/2 that disrupts MCM5 ATPase activity required for DNA replication initiation, produced a comparable response by significantly reducing viability in shCON LNCaP cells and produced even stronger inhibition in shCRY1 cells (**Fig. 5E**). Notably, co-treatment with either APLF or MCM5 inhibitors with a CRY1 inhibitor recapitulated the effects observed in CRY1 knockdown models, indicating an additive effect on suppression of cell viability (**Fig. 5F**). Together, these findings indicate that pharmacologic inhibition of CRY1 phenocopies genetic depletion, reinforcing a functional link between CRY1 activity and replication-associated vulnerabilities and supporting CRY1 as a tractable therapeutic target in prostate cancer Treatment-response differences were evaluated using two-way ANOVA with multiple-comparison correction. Although refametinib is best characterized as a selective MEK1/2 inhibitor, its capacity to intensify replication stress likely underlies the MCM-linked vulnerability observed here. In contrast, CRPC-identified inhibitors showed minimal activity in LNCaP (HTS), and co-treatment with a CRY1 inhibitor did not enhance responses in parental LNCaP cells (**Supplementary Fig. 5A-B**), indicating HTS-restricted dependence on APLF/MCM-associated processes.

Importantly, in C4-2 (CRPC) a setting where HR programs predominate CRY1 loss exposed dependencies on ATM and ERCC3. The ATM inhibitor, KU-55933 (2.5 µM), produced moderate inhibition in shCON cells, with substantially enhanced sensitivity in shCRY1 cells. Similarly, BPK-21 (5 µM), a covalent ERCC3/XPB inhibitor, recapitulated this pattern (**Fig. 5C-D**). In parental C4-2 cells, dual inhibition of CRY1 and ATM or ERCC3 further suppressed viability (**Fig. 5G-H**). Treatment-response differences were evaluated using two-way ANOVA with multiple-comparison correction.

These results suggest that co-targeting CRY1 and key DNA damage response pathways represents a rational combination strategy to enhance therapeutic vulnerability in PCa. By potentiating the effects of PARP, APLF, MCM5, ATM, and ERCC3 inhibitors, CRY1 inhibition may amplify unrepaired DNA damage, either at NHEJ in the early stage of the disease or at HR as the stage of the disease progresses and drives tumor cell lethality. Notably, early implementation of this approach may delay progression to castration-resistant disease, highlighting its potential to both improve durability of response and shift the treatment paradigm toward prevention of advanced PCa.

By contrast, HTS specific inhibitors showed a minimal effect in C4-2 cells, and their activity was not significantly improved by CRY1 inhibition (**Supplementary Fig. 5C-D**). The role of CRY1 in directing HR programs in PCa was elegantly defined in our previous study^7^, which demonstrated that CRY1 governs multiple tiers of the HR machinery including upstream sensors, key mediators, and downstream effectors thereby positioning CRY1 as a central transcriptional regulator of HR fidelity in advanced PCa^7^. Together, these data highlight CRPC-specific vulnerabilities to ATM/ERCC3 blockade.

To further define combinatorial vulnerabilities, we assessed the interaction between CRY1 inhibition and DDR-targeted agents in both HTS (LNCaP) and CRPC (C4-2) models using the highest single agent (HSA) model for synergy assessment. In LNCaP cells, co-treatment with PARP inhibition demonstrated strong synergy (mean HSA score=12.7), indicating robust enhancement of therapeutic efficacy upon CRY1 targeting. A similar degree of synergy was observed in C4-2 cells treated with CRY1 inhibitor and PARP inhibitor (mean HSA score=12.3), suggesting that PARP dependency remains sensitized by CRY1 inhibition across disease states. In contrast, combinations targeting alternative DDR pathways showed more context-dependent effects. In LNCaP cells, CRY1 inhibition combined with APLF yielded modest synergy (mean HSA score=9.25), approaching but not exceeding the defined synergy threshold. Notably, in CRPC cells, co-targeting of CRY1 and ATM resulted in strong synergistic interactions (mean HSA score=12.3), highlighting a pronounced dependence on ATM-mediated repair pathways in advanced disease (**Supplementary Fig.5E**).

Collectively, these findings indicate that CRY1 inhibition enhances sensitivity to DDR-targeted therapies in a pathway- and context-dependent manner, with consistent PARP-associated synergy across both HTS and CRPC models and pronounced ATM-directed vulnerability in CRPC. More broadly, these data reveal that loss of CRY1 unmasks distinct DDR liabilities across prostate cancer states, characterized by APLF/MCM-linked dependencies in HTS and ATM/ERCC3 dependencies in CRPC. This bifurcation is consistent with a transition from NHEJ-biased repair in HTS to HR-dominant programs in CRPC.

Within this framework, CRY1 appears to function as a dynamic regulator of repair pathway engagement, integrating proliferative and stress signals to coordinate DDR program selection across disease progression. Consistent with prior observations, CRY1 is stabilized following DNA damage and contributes to temporal regulation of HR-associated gene expression. In this context, PARP activity likely represents a critical intermediate state, reinforcing reliance on PARP-mediated BER under conditions of replication stress. Together, these results support a model in which CRY1 sustains genomic integrity through adaptive DDR regulation, and its loss exposes stage-specific repair vulnerabilities that may be therapeutically exploited.

### CRY1 Co-expression and Genomic Co-alteration with DDR Genes Associate with Adverse Clinical Outcomes in PCa

To elucidate the potential clinical relevance of our pre-clinical findings, we examined publicly available datasets of human PCa samples. Co-expression analysis of CRY1 with the validated mRNA targets *PARP1, APLF, ATM*, and *ERCC3* mRNA was further assessed across multiple independent cBioPortal cohorts, including MSK, Cancer Cell 2020, DKFZ, Cancer Cell 2020, MSK, and SU2C/ PCF Dream Team, Cell 2015, and revealed significant positive correlations. In the primary cohort (**Fig. 6A**), *CRY1* expression correlated with *PARP1* (p value = 4.95^e-7^), *APLF* (p value = 2.52^e-5^), *ATM* (p value = 2.70^e-12^), and *ERCC3* (p value = 2.13^e-4^). These relationships were reproducible across independent cohorts (**Supplementary Fig. 6A**), including additional datasets, SU2C/ PCF Dream Team, PNAS 2019, SU2C/ PCF Dream Team, Cell 2015, showing consistent correlations (for example, *PARP1*: p value = 7.066^e-4^ and 3.030^e-3^; *ATM*: p value = 0.014 and 1.26^e-7^; *ERCC3*: p value = 3.310^e-3^), highlighting a stable and clinically relevant transcriptional link between CRY1 and key DNA damage response components.

**Figure 6.**
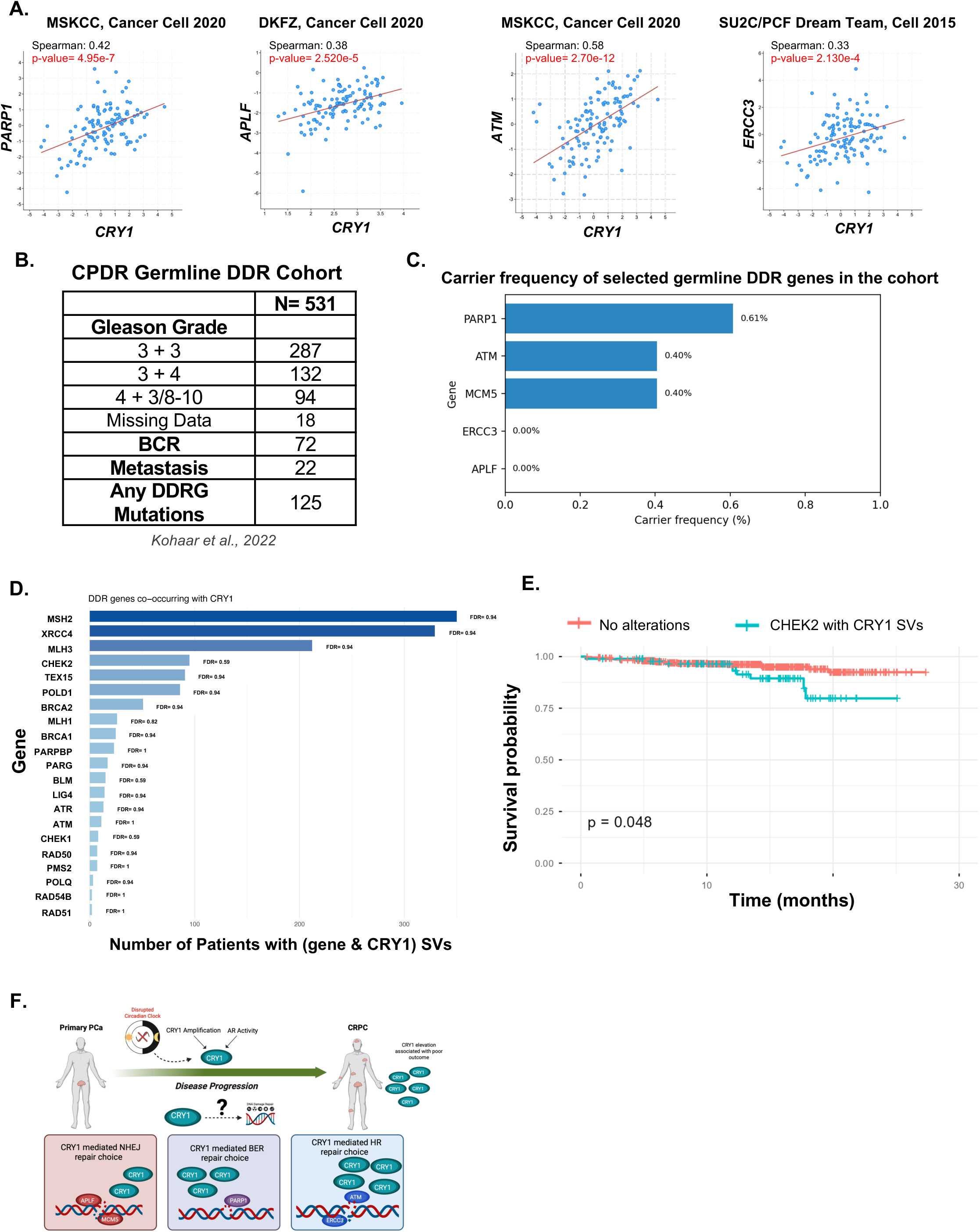
**A.** Co-expression analysis of CRY1 with PARP1, APLF, ATM, or ERCC3 mRNA in prostate cancer (PCa) tissues. Analyses were performed using publicly available datasets, including MSKCC (Cancer Cell, 2020), DKFZ (Cancer Cell, 2020), and the SU2C/PCF Dream Team cohort (Cell, 2015). **B.** Summary of germline DDR alterations in the CPDR cohort (N=531), including Gleason grade, BCR, metastasis, and DDR mutation frequency (adapted from *Kohaar et al., Nat. Commun.*, 2022). **C.** Single-nucleotide variant (SNV) carrier frequencies of selected DNA damage repair drug targets in the CPDR cohort (n = 531). Analysis was restricted to dnSNV carriers, with CHEK2 excluded to avoid confounding effects from its high prevalence. **D.** Frequency of DDR structural variants co-occurring with CRY1 structural variants in the CPDR cohort (n = 531), highlighting genes involved in alternative DNA damage repair pathways. **E.** Kaplan–Meier survival analysis comparing patients with co-occurring CRY1 and CHEK2 structural variants versus patients without these alterations in the CPDR cohort (n = 531). Statistical significance was assessed using the log-rank test. **F.** Model summarizing the findings from this study, illustrating CRY1-associated disease progression and a proposed shift in DNA repair pathway choice driven by compensatory activation of alternative DDR mechanisms.

Moreover, the relationship between *CRY1* and DDR alterations was examined in a previously published large germline DDR PCa cohort (n = 531) (**Fig. 6B**)^38^. Within this cohort, pathogenic or likely pathogenic variants were identified across multiple DDR genes, with measurable carrier frequencies observed for canonical repair factors including *PARP1* and *ATM* (**Fig. 6C**). Notably, structural variations involving *CRY1* co-occurred with alterations in a broad spectrum of DDR genes, most prominently *MSH2, XRCC4, MLH3*, and *CHEK2* (**Fig. 6D**), indicating a non-random overlap between *CRY1* genomic alteration and compromised genome maintenance pathways. Clinically, patients harboring *CHEK2* alterations alongside *CRY1* structural variants experienced significantly reduced survival compared with those lacking these alterations (p = 0.048) (**Fig. 6E**). This is consistent with the role of CHEK2 as a key checkpoint kinase downstream of ATM that regulates cell cycle arrest and facilitates homologous recombination through BRCA1-dependent mechanisms^39^.

Together, these observations support a model in which CRY1 alterations intersect with inherited defects in DNA repair, further implicating CRY1 as a contributor to aggressive disease biology and adverse clinical outcomes in PCa. In summary, our data support a model in which CRY1 functions as a context-dependent protumorigenic factor that becomes amplified as PCa progresses, shaping DNA repair pathway utilization across disease states. In HTS, CRY1 preferentially utilizes NHEJ, whereas with disease progression it facilitates a shift toward HR in CRPC, with an intermediate role in BER. Through dynamic regulation of key repair factors across these pathways, CRY1 acts as a central determinant of DNA repair plasticity, enabling tumor adaptation and progression (**Fig. 6F**).

## Discussion

Disruption of circadian regulation has long been linked to cancer risk and progression, yet how circadian factors interface with genome maintenance programs during tumor evolution has remained incompletely understood. Core circadian components, including cryptochromes, have been implicated in coordinating cellular responses to genotoxic stress and regulating DNA damage-induced apoptosis, highlighting a functional connection between circadian timing and DDR pathways^5,7^. In this study, we identify the core circadian clock regulator CRY1 as a stage-contingent organizer of DDR pathway utilization in PCa. Building upon our prior work establishing CRY1 as a pro-tumorigenic factor that promotes HR in advanced disease^7^, the present findings demonstrate that CRY1 dynamically rewires DNA repair circuitry as tumors progress from HTS to CRPC. By coupling proliferative cues to genome maintenance pathways, CRY1 enables tumor cells to tolerate genotoxic stress during disease progression (**Fig. 2 C-D**). Disruption of this coupling through CRY1 loss exposes a vulnerability in PCa cells, particularly in advanced disease states^7,40^.

By integrating transcriptomic profiling, focused DDR CRISPR screening, pharmacologic perturbation, and clinical correlative analyses, we show that CRY1 loss does not uniformly compromise genome maintenance but instead unmasks distinct, disease-state specific repair dependencies. In HTS models, CRY1 preferentially aligns with BER and NER associated transcriptional and functional programs, consistent with reliance on repair mechanisms that resolve single-strand lesions and replication-associated stress in early disease. In contrast, CRPC models exhibit pronounced CRY1-dependent engagement of HR-associated pathways, reflecting the heightened need for high-fidelity double-strand break repair under conditions of augmented genomic instability and therapeutic pressure (**Fig. 6F**). These findings are consistent with prior studies demonstrating that advanced PCa is enriched for alterations in HR and other DDR pathways, including BRCA1/2 and ATM, which are frequently observed in metastatic CRPC^41^. Notably, circadian regulation of DNA repair pathway choice has been linked to CRY1-dependent modulation of DNA end resection, a key determinant of HR versus NHEJ utilization, providing a potential mechanistic basis for the stage-specific rewiring observed here^42^.

The enrichment of MMR signatures within the overlapping CRY1-regulated gene set suggests that PCa cells engage intermediate repair strategies during disease evolution, consistent with adaptive DDR pathway switching models. Such plasticity may enable tumors to buffer accumulating genomic insults while maintaining sufficient replicative capacity to support progression. Within this framework, CRY1 appears to function less as a static DDR regulator and more as a contextual transcriptional hub that coordinates repair pathway selection in accordance with disease stage and cellular stress.

An important revelation from this study is the emergence of MMR as a shared, transitional program bridging HTS and CRPC states. Given its role in correcting replication-associated mismatches, MMR may provide a selective advantage during the transition to castration resistance by mitigating replication stress without requiring fully engaged double-strand break repair. In this context, MMR activity could stabilize replication forks and limit accumulation of deleterious mutations, thereby preserving cellular viability under increasing proliferative and therapeutic pressure. Thus, CRY1-associated induction of MMR programs may function as an adaptive safeguarding mechanism that enables tumor cells to tolerate genomic stress while maintaining the capacity for continued progression^43,44^.

In practical terms, this framework provides a rationale for precision combinations that leverage CRY1 inhibition with pathway-matched partners and BER liabilities in HTS and HR vulnerabilities in CRPC while also prioritizing common transitional nodes (e.g., MMR) for interception strategies. Such stage-informed targeting could enable more durable responses by aligning therapeutic pressure with the dominant, CRY1-organized repair axis operating in each disease state.

These transcriptional programs are further supported by Hallmark pathway analysis, which reveals that CRY1-regulated gene networks evolve with disease progression. In HTS models (LNCaP), enriched pathways are associated with proliferation and cellular homeostasis, consistent with androgen-driven growth. Genes shared between HTS and CRPC converge on cell cycle and DDR-associated programs, whereas CRPC-specific gene sets (C4-2) are enriched for inflammatory signaling, cytokine pathways, and stress-responsive networks, hallmarks of castration-resistant disease biology. These observations align with prior studies demonstrating that CRPC is characterized by transcriptional reprogramming toward stress adaptation and therapeutic resistance, often accompanied by enhanced DDR activity and reliance on replication stress response pathways^45^.

The functional consequences of this rewiring are underscored by our CRISPR-based dependency mapping and pharmacologic validation (**Figs. 3-5**). CRY1 depletion preferentially sensitized HTS models to disruption of APLF- and MCM5-associated processes, consistent with reliance on replication-associated repair and chromatin remodeling, while revealing CRPC-specific vulnerabilities to ATM and ERCC3 inhibition. In contrast, CRPC models exhibited heightened sensitivity to inhibition of ATM and ERCC3, key regulators of HR-mediated repair and transcription-coupled nucleotide excision repair, respectively. These findings highlight a partitioning of DDR liabilities that mirrors transcriptional reprogramming during disease progression. PARP1 emerged as a shared dependency across disease states, consistent with its central role in BER, replication fork stability, and coordination of DNA repair signaling networks. Mechanistically, PARP inhibition induces accumulation of DNA lesions and increases reliance on ATR-CHK1 mediated replication stress responses, providing a rationale for combinatorial targeting strategies^46^.

While our analysis is not directly addressed in prior studies, recent work provides relevant conceptual context for interpreting trends in CRISPR-edited genomic data. It has been shown that disruption of DDR pathways can allow cells to persist despite accumulating genomic damage, particularly under selective pressure, without necessarily producing strong binary phenotypes^47^. It has also been demonstrated that DDR genes often function within highly interconnected networks where individual gene effects are modest and context-dependent, emerging as gradual trends rather than sharp on/off signals^48^. Together, these studies support interpreting our observed association as a progression-linked pattern of cumulative genomic disruption, rather than evidence of a discrete driver event.

Importantly, pharmacologic inhibition experiments demonstrated that CRY1 loss or inhibition enhances sensitivity to DDR inhibitors in a manner consistent with these stage-specific dependencies. In HTS models, CRY1 disruption amplified responses to inhibitors impacting replication-associated and BER-linked processes. In this context, elevated CRY1 expression may identify tumors with intact but stress-adapted repair capacity, for which CRY1 inhibition could unmask vulnerabilities to agents that exacerbate replication stress or disrupt BER/NER-linked processes. This raises the possibility that CRY1 status could stratify patients who are less likely to benefit from HR-directed therapies but may instead respond to alternative DDR-targeted combinations earlier in the disease course.

In CRPC models, CRY1 inhibition markedly increased susceptibility to ATM and ERCC3 blockade nodes closely tied to HR integrity and transcription-coupled repair. CRY1 inhibition may functionally compromise HR capacity and thereby expand the population of CRPC patients who could derive benefit from PARP inhibitors or other HR-disrupting agents, independent of BRCA mutation status. These results suggest that CRY1 inhibition effectively lowers the buffering capacity of DDR networks, exposing latent vulnerabilities that can be therapeutically exploited when paired with pathway-matched DDR inhibitors.

The stage-contingent role of CRY1 in directing DDR pathway choice has important implications for patient stratification and therapeutic decision-making in PCa. Current clinical approaches to DDR-targeted therapy, including PARP inhibition, predominantly rely on static genomic markers such as BRCA1/2 or other HR repair gene alterations. However, these biomarkers incompletely capture the dynamic rewiring of repair dependencies that accompanies disease progression and therapeutic pressure. Our findings suggest that CRY1 expression and activity may provide complementary, functional information that reflects the prevailing DDR state of a tumor rather than merely its mutational history.

Importantly, the identification of lineage-restricted DDR dependencies downstream of CRY1 loss suggests a framework for stage-aware combination therapies. In HTS disease, CRY1 inhibition combined with agents targeting APLF or MCM-associated pathways may selectively impair tumor growth while sparing normal tissues. In contrast, CRPC tumors characterized by CRY1-driven HR reliance may be particularly susceptible to combinations of CRY1 inhibitors with ATM or ERCC3 blockade. Such stratified approaches could help align therapeutic pressure with the dominant repair axis operating in each disease state, potentially improving durability of response and delaying resistance. Notably, inhibition of HR pathways can sensitize tumors to PARP inhibitors, and combination strategies targeting ATR or CHK1 further exacerbate replication stress and promote tumor cell death. These findings suggest that CRY1 inhibition lowers the buffering capacity of DDR networks, exposing synthetic vulnerabilities that can be exploited through rational therapeutic combinations^49^. Our findings suggest that CRY1 expression and activity may provide complementary, functional insight into the prevailing DDR state of a tumor.

Our clinical analyses further support the relevance of CRY1-associated DDR circuitry in human disease (Fig. 6). CRY1 dysregulation correlated with poor clinical outcome, co-expression with critical DDR genes, and non-random co-occurrence with inherited or somatic DDR alterations. Notably, patients harboring concurrent CRY1 and CHEK2 structural variants exhibited significantly reduced survival (Fig. 6E), suggesting that CRY1 may cooperate with compromised checkpoint signaling to accelerate disease aggressiveness. While additional studies will be required to delineate causality, these observations position CRY1 at the intersection of circadian regulation, DDR integrity, and clinical progression. Notably, concurrent alterations in *CRY1* and checkpoint regulators such as *CHEK2* may further exacerbate genomic instability and disease aggressiveness, consistent with the broader observation that DDR pathway disruptions contribute to advanced disease phenotypes and poor prognosis in PCa ^44^.

Thus, this work advances a model in which CRY1 functions as a temporal gatekeeper of genome maintenance, dynamically steering DNA repair pathway choice as PCa evolves ( **Fig. 6F**). This stage-aware conceptual framework has direct therapeutic implications. Rather than uniformly targeting DDR pathways, our data support precision strategies that combine CRY1 inhibition with BER/NER-focused interventions in early disease and HR-directed approaches in advanced CRPC. More broadly, these findings highlight circadian regulators as active architects of DDR plasticity, whose dysregulation contributes to genomic adaptation, therapeutic resistance, and cancer progression.

In summary, these findings position CRY1 as a potential dynamic biomarker of DDR state that integrates circadian regulation, transcriptional control, and genome maintenance. Incorporating CRY1 expression or activity into clinical stratification frameworks may enable more precise matching of patients to DDR-targeted therapies across disease stages. Future prospective studies integrating CRY1 profiling with longitudinal sampling and therapeutic response data will be essential to evaluate whether CRY1-guided strategies can improve outcomes beyond current genomics-based approaches.

## Materials and Methods

### Cell Culture and Reagents

The LNCaP and C4-2 PCa cell lines were purchased from the American Type Culture Collection (ATCC), and they provided initial authentication. Mycoplasma testing (Cat#30-1012K) was performed to ensure all cells were negative for the experiments performed in this study. Following thawing, all cells were screened for mycoplasma contamination by PCR. Cells were maintained in RPMI medium (Corning, 10-040-CV) supplemented with 5% heat-inactivated fetal bovine serum (FBS), 2 mmol/l L-glutamine, and 100 units/ml penicillin-streptomycin. Cultures were incubated under standard conditions at 37°C in a 5% CO_2_ environment. Doxycycline-inducible CRY1 knockdown models (shCRY1 and shCON control) were generated as previously detailed^7^.

### RNA Isolation and Sequencing

LNCaP cells (ATCC) were authenticated upon receipt and confirmed mycoplasma-free prior to use. After experimental treatments, total RNA was extracted using the miRNeasy kit (Qiagen) following the manufacturer’s protocol. RNA integrity was confirmed using an Agilent Bioanalyzer, and only samples with RIN>8 were processed further. RNA-seq library construction was performed at the Sidney Kimmel Cancer Center Sequencing Core Facility using the TruSeq Stranded Total RNA Library Prep Gold kit (Illumina). Libraries were sequenced on an Illumina NextSeq 500 using single-end 75 bp reads.

### RNA-sequencing Analysis

FASTQ files were assessed via FASTQC, and reads were aligned to the human genome (hg19) with STAR v2.5.2a. Read quantification was performed with featureCounts using Ensembl gene annotation. Prior to downstream processing, raw count matrices underwent an initial cleaning step in which entries labeled as NA (genes lacking valid identifiers or mapping annotation) were removed to ensure accurate and biologically interpretable quantification. For comparison between the LNCaP and C4-2 datasets, additional filtering was implemented to harmonize detected gene sets. Specifically, the union of expressed genes across both datasets was examined, and only genes meeting a statistical significance threshold of p<0.01 in differential expression testing were retained. This filtering was conducted prior to pathway analysis to minimize noise and restrict analysis to high-confidence, biologically relevant transcripts. Following filtration, differential gene expression analysis was performed using DESeq2 with shrinkage estimation for dispersion and fold change. Gene set enrichment analysis (GSEA) was performed using KEGG and HALLMARK gene signatures.

### Quality Control and Filtering

To ensure comparability with the previously published C4-2 RNA-seq dataset from Shafi *et al*.^7^, identical quality control principles were applied. Sample-level QC included principal component analysis, replicate clustering, and evaluation of alignment metrics, duplication rates, and library complexity. Filtering of NA-annotated genes and retention of only statistically significant genes (p<0.01) prior to pathway analysis ensured that downstream comparisons between LNCaP and C4-2 were conducted using harmonized, high-confidence gene sets.

### Immunoblotting

Cells were plated at uniform densities in hormone-proficient media. Cell lysates were prepared as previously described^50^, and 20–50 µg of total protein was resolved via SDS-PAGE. Proteins were transferred to PVDF (polyvinylidene fluoride) membranes and probed with primary antibodies at 1:1000 and 1:5000 dilutions for CRY1 and Vinculin respectively. Specific antibodies included CRY1 (Bethyl A302-614A) and Vinculin (Sigma–Aldrich V9264), the latter serving as a loading control.

### Cell Viability Assays

To assess the impact of targeted inhibition on cell viability, LNCaP (2,000 cells/well) and C4 -2 (1,000 cells/well) were seeded into white-walled, clear-bottom 96-well plates. After a 24-hour attachment period, cells were treated in triplicate for 72 hours with a panel of inhibitors from MedChemExpress (MCE), including CRY1 inhibitor (M47, HY-148764), APLF inhibitor (ETP-46464, HY-15521), MCM5 inhibitor (Refametinib, HY-14691), ATM inhibitor (KU-55933, HY-12016), and ERCC3 inhibitor (BPK-21, HY-141549). Cell viability was determined using the CellTiter-Glo 2.0 (Promega #G9242) luminescent readout, with all experimental groups normalized to DMSO vehicle controls.

### Xenograft Analysis

C4-2 PCa cells harboring doxycycline-inducible shRNA constructs targeting CRY1 (C4-2-shCRY1) or control (C4-2-shCON) were resuspended in 100 μL of a 1:1 mixture of Matrigel (50%, BD Biosciences) and sterile saline and subcutaneously injected into male athymic nude mice (≥6 weeks old). Tumor growth was monitored longitudinally, and tumor volumes were calculated using caliper measurements. Following tumor establishment, mice were administered doxycycline-supplemented drinking water (2 mg/mL doxycycline with 5% sucrose) to induce CRY1 knockdown. Treatment continued for the duration of the experiment, and tumor growth was assessed over time. Mice were sacrificed when tumors reached endpoint criteria (tumor volume ≥1000 mm³), and tumors were harvested for downstream analyses. All animals used in this study were male, and no mice experienced greater than 5% body weight loss during the study period. All procedures were approved by the Institutional Animal Care and Use Committee (IACUC) and conducted in accordance with institutional guidelines and all relevant regulations for the ethical use of animals in research.

### PDE (patient derived explant)

Fresh PCa specimens were obtained with written informed consent from men undergoing robotic radical prostatectomy at Walter Reed National Military Medical Center. Dissected tissue fragments were utilized as *ex vivo* PDE cultures as previously described ^16,51-53^.Institutional Review Board has reviewed this protocol and deemed this research to follow federal regulations (Approval #DBS.2024.699). PDE cultures were treated with media containing M47 (100 and 150 mM) or vehicle alone (DMSO) and harvested after 72 hours for hematoxylin and eosin (H&E) and immunohistochemistry (IHC) analyses. For histological analysis, explants were formalin-fixed and paraffin embedded (FFPE). H&E staining was performed using standard protocols following standard procedures in a CLIA-certified clinical laboratory at HistoServ (Germantown, Maryland, USA). Glass slides were then scanned and digitalized images used for further assessment.

### CRISPR DDR Library Design and Construction

A focused human DNA damage response (DDR) CRISPR Cas9 sgRNA library was used to target 365 genes with established or putative roles in DNA repair and genome maintenance, supplemented with 45 core essential genes and 50 nonessential genes as internal controls. The DDR gene set incorporated 275 genes described and expanded based on previous studies^23,54^ and further refined through expert curation of additional DDR-related pathways. Where feasible, 10 sgRNAs were designed per gene, resulting in a total of 4,530 sgRNAs.

### Lentivirus Production

Lentiviral particles were generated in 293FT cells using a second-generation packaging system. Cells were seeded one day prior to transfection to achieve ∼60% confluence. For each 15-cm dish, cells were transfected with lentiCRISPRv2 DDR library plasmid (20 µg), psPAX2 (14 µg), and PMD2.G (9 µg) using TransIT-Mirus reagent (3:1 reagent-to-DNA ratio) in Opti-MEM. Viral supernatants were collected at 72 and 96 hours post-transfection, clarified by centrifugation, and concentrated by PEG precipitation. Viral pellets were resuspended in PBS and stored at −80 °C or used immediately.

### Pooled CRISPR Screening

LNCaP shCON, LNCaP shCRY1, C4-2 shCON, and C4-2 shCRY1 Doxycycline inducible cell lines were transduced with the DDR CRISPR library at a multiplicity of infection (MOI) of ∼0.3 to ensure single integrants, while maintaining a minimum 1,000X coverage per sgRNA. Following transduction, cells were selected with puromycin for 24 hours and then maintained induced with 1 µg/ml of doxycycline, replenishing doxycycline every 72 hours. A baseline reference sample (T0) was collected immediately after selection. Cells were subsequently cultured for up to 10 passages. Cell pellets were collected longitudinally for genomic DNA and protein analyses. Library maintenance was confirmed by PCR detection of sgRNA cassettes, demonstrating robust retention of viral constructs at early (P0+V) and late (P10+V) passages, with no detectable signal in non-transduced controls (P0−V).

### Genomic DNA Extraction and Sequencing

Genomic DNA was isolated from baseline (T0) and endpoint (T21) samples using the QIAamp DNA Blood Midi Kit (Qiagen) according to the manufacturer’s instructions. sgRNA cassettes were amplified from genomic DNA by a two-step PCR approach. In the first PCR, sgRNA regions were amplified using high-fidelity DNA polymerase to ensure accurate representation of guide sequences. A second PCR was performed to append sequencing adapters and sample-specific indices using the following primers: hU6-F (5′-GAG GGC CTA TTT CCC ATG ATT-3′) and LentiCRISPR_v2_sgRNA-scaffold_rev (5′-CCT TAT TTT AAC TTG CTA TTT CTA GCT CTA AAA C-3′).

PCR products (∼350 bp) were size-verified by agarose gel electrophoresis and purified using AMPure XP beads (Beckman Coulter). Libraries were quantified, pooled, and sequenced on an Illumina NextSeq 500 platform using 75-bp single-end reads.

### CRISPR Data Processing and Analysis

Sequencing data from pooled CRISPR–Cas9 screens were processed using a customized analysis pipeline optimized for the specific sgRNA library architecture used in this study. Initial attempts to align sequencing reads using the standard MAGeCK-VISPR alignment workflow were unsuccessful due to library-specific sequence features that interfered with automated sgRNA identification. As a result, a deconstructed MAGeCK-based pipeline was implemented to ensure accurate sgRNA quantification and downstream analysis.

Raw FASTQ files were first subjected to preprocessing using trimming procedures, in which constant adapter and flanking regions inherent to the sgRNA library design were removed to isolate the 20 bp sgRNA targeting sequence. Specifically, fixed-length sequences were trimmed from the 5′ (35 bp) and 3′ (54 bp) ends of each read, corresponding to known repetitive and non-informative regions. This trimming step was critical for standardizing read length and preventing misalignment due to residual library backbone sequence.

Following trimming, the sgRNA reference library was converted from a tabular (CSV) format into FASTA, and index files were generated using the Burrows–Wheeler Aligner (BWA). Trimmed reads were then aligned to the sgRNA reference using BWA, enabling precise and reproducible mapping of sequencing reads to individual sgRNAs. This explicit mapping strategy ensured accurate handling of the library’s structure and avoided alignment artifacts observed with automated MAGeCK alignment routines.

Aligned reads were quantified using the mageck count function with BWA-aligned reads as input. Internal quality control metrics were assessed at this stage, including sgRNA representation, read depth distribution, and consistency across replicates. Read counts were median-normalized across samples to account for differences in sequencing depth.

Normalized count tables were combined across conditions and biological replicates for downstream statistical analysis. Differential sgRNA abundance was evaluated using both the MAGeCK test and MAGeCK maximum likelihood estimation frameworks to identify genes exhibiting consistent enrichment or depletion across multiple sgRNAs. Final outputs included ranked gene lists, gene-level summary statistics, and sgRNA-level summaries.

All downstream statistical analyses and visualizations were performed in R, including alternative enrichment plots and comparative diagrams to facilitate interpretation of screen performance and candidate gene prioritization. While the standard MAGeCK-VISPR pipeline may be suitable for other experimental contexts, the customized trimming and BWA-based alignment workflow described here was developed to ensure robustness and accuracy for the sgRNA library used in this study.

### Co-expression Analysis

We queried cBioPortal to obtain gene correlation metrics derived from the indicated studies^55,56^.

## Acknowledgements

This work was supported by a Young Investigator Award and Challenge Award from the Prostate Cancer Foundation (to A.A.S., A.F., and K.E.K.), NCI R01-CA182569 (K.E.K.), the Sidney Kimmel Cancer Center (SKCC) Support Grant (P30CA056036), CPDR-CORE funds (A.A.S., X.A.S.), American Cancer Society (ACS) Postdoctoral Fellowship (PF-24-1318851) (F.P.) in addition to the Cancer Genomics and Translational Research and Pathology Core Services at SKCC. The collaborators contributed to the design of the study, data collection, analysis, interpretation, and review and approval of the manuscript. We are grateful for their valuable insights and support throughout this work. We also thank former members of the Knudsen laboratory and current CPDR colleagues for constructive discussions and technical assistance.

## Ethical approval

The use of human specimens and associated clinical data in this study was conducted under Institutional Review Board (IRB)–approved protocols at the Uniformed Services University (USU) and the Center for Prostate Disease Research (CPDR) (protocol: BCBR, DBS.2024.699). Patient-derived explants (PDEs) were generated from PCa tissue obtained from patients undergoing surgery with appropriate informed consent. Additional patient clinical and genomic data were obtained from the CPDR cohort in accordance with approved protocols governing the collection, storage, and analysis of human specimens. All studies involving human tissues were performed in compliance with institutional guidelines and in accordance with all applicable ethical regulations for research involving human subjects.

## Author contributions

AF, AAS conceived and designed the project. AF, SD, LR, TJ, FP, LS, NEA, OR, SE, AA, JJ, IS, CS ML, GS, XAS, CMM, KEK, AAS acquired the data. AF, SD, FP, CMM, and AAS analyzed and interpreted the data. AF, SD, and AAS wrote the paper.

