## Supplemental Figures for "Stage-Specific Regulation of DNA Damage Repair by the Circadian Regulator, CRY1, in Prostate Cancer": Fallatah et al. 2026_Figures - Supplemental.pdf

### Supplementary Figure 1

A.

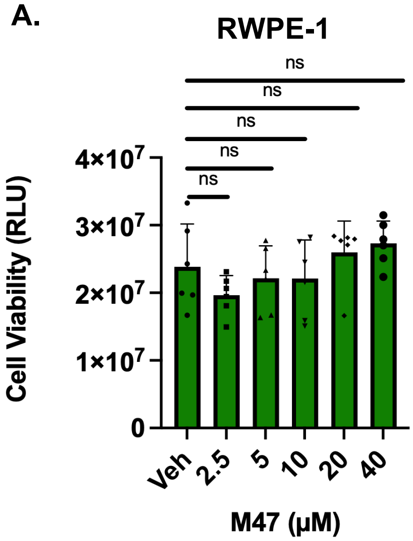


B.

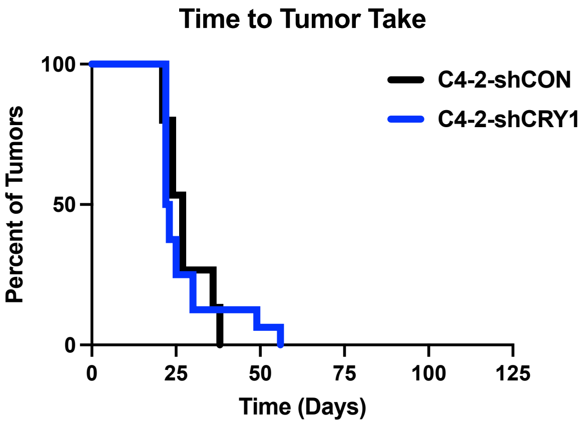


Supplementary Figure 1. **CRY1 is Protumorigenic and Inhibition Decreases Growth in PCa. A.** Cell Titer-Glo 2.0 Cell Viability Assay in RWPE-1 (normal prostate epithelial cell line) upon treatment with CRY1 inhibitor (M47). ns= not significant. **B.** Time it takes the tumor to develop in C4-2 shCON and C4-2 shCRY1.

### Supplementary Figure 2

**A.** LNCaP PCA

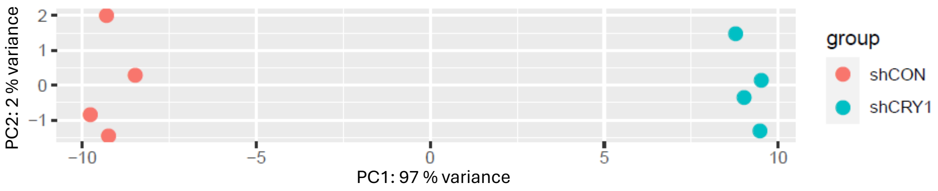


**B.** LNCaP Unique Genes GSEA (*Hallmark*)

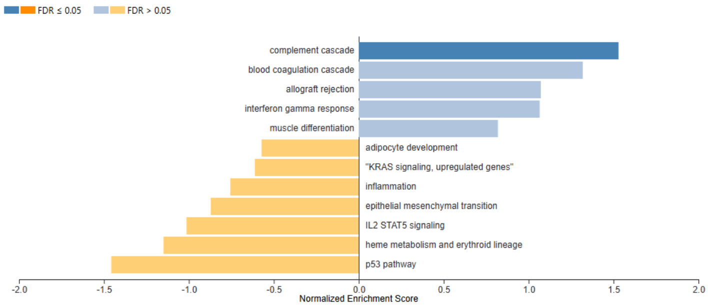


**C.** LNCaP and C4-2 Common Genes GSEA (*Hallmark*)

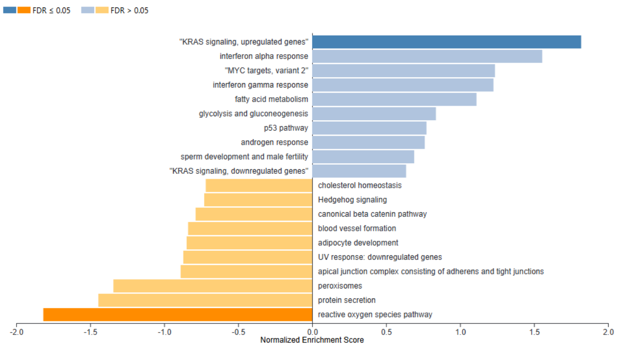


**D.** C4-2 Unique Genes GSEA (*Hallmark*)

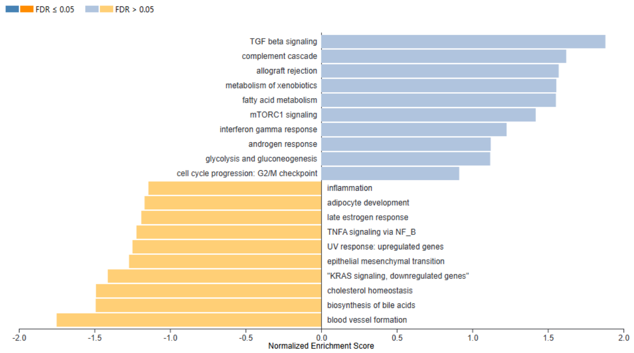


<u>Supplementary Figure 2.</u> **CRY1 Transcriptional Landscape in HTS and CRPC Models. A.** PCA plot of LNCaP shCON and shCRY1 cells. **B-D.** GSEA Hallmark pathway enrichment analysis of differentially expressed genes in LNCaP unique (B), common to LNCaP and C4-2 (C), and C4-2 unique (D).

### Supplementary Figure 3

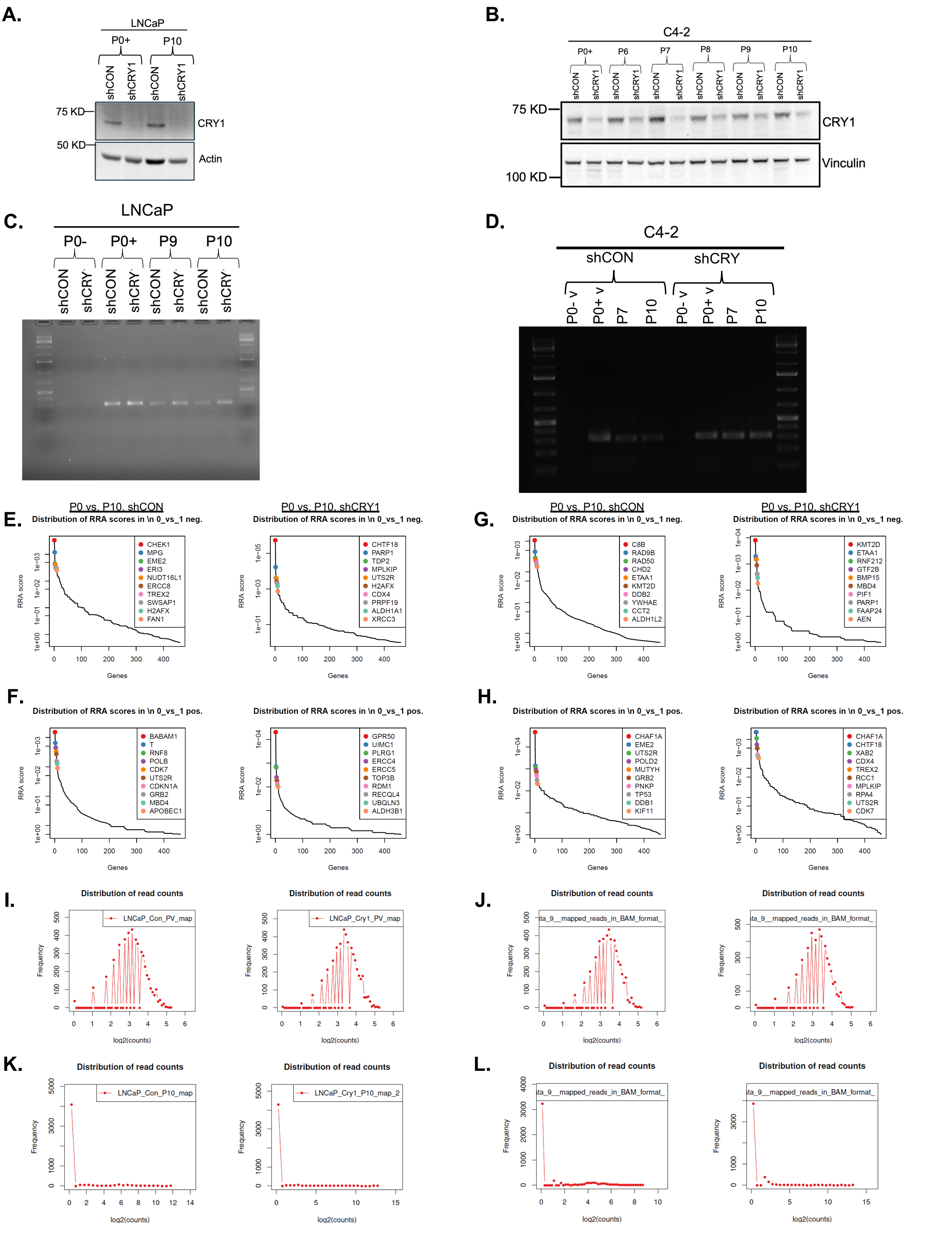

### Supplementary Figure 3

**Supplemental Figure 3. DNA Damage Repair CRISPR screen with CRY1 knock down. (A-B).** Western blot analysis confirms stable CRY1 knockdown across all passages in LNCaP (**A**) and C4-2 (**B**) cells. (**C,D**) Agarose gel electrophoresis showing the results of the DDR CRISPR library screen performed in LNCaP (HTS; **C**) and C4-2 (CRPC; **D**) disease models following CRY1 knockdown via shRNA. **E.** Ranked Robust Rank Aggregation (RRA) curve, LNCaP, shCON, CRY1, negative selection. **F.** Ranked Robust Rank Aggregation (RRA) curve, LNCaP, shCON, CRY1, positive selection. **G.** Ranked Robust Rank Aggregation (RRA) curve, C4-2, shCON, CRY1, negative selection. **H.** Ranked Robust Rank Aggregation (RRA) curve, C4-2, shCON, CRY1, positive selection. **I.** Distributions of sgRNA read counts across biological replicates, LNCaP, P0. **J.** Distributions of sgRNA read counts across biological replicates, C4-2, P0. **K.** Distributions of sgRNA read counts across biological replicates, LNCaP, P10. **L.** Distributions of sgRNA read counts across biological replicates, C4-2, P10.

### Supplementary Figure 4

**A.** **LNCaP shCRY1** MLE

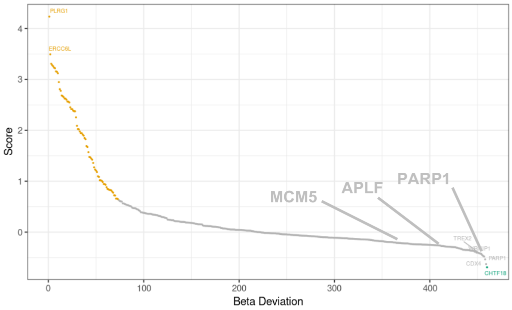


**B.** **C4-2 shCRY1** MLE

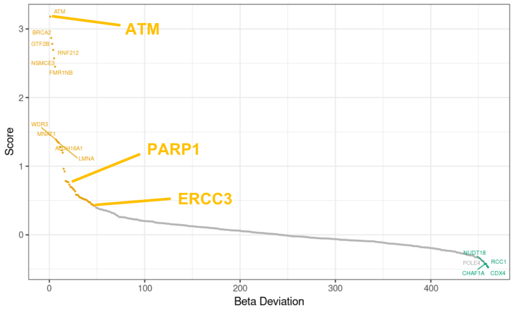


**C.**

**HALLMARK Pathways for LNCaP** negative RRA selections (p< 0.05)

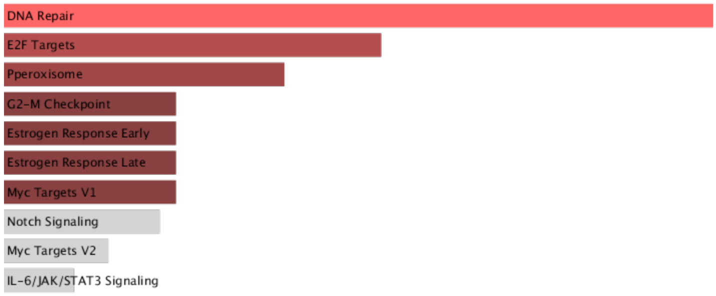


**D.**

**HALLMARK Pathway for Common** negative RRA selections (p< 0.05)

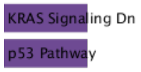


**E.**

**HALLMARK Pathway for C4-2** negative RRA selections (p< 0.05)

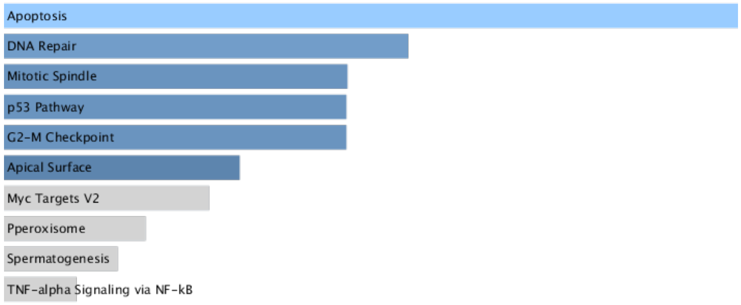


**F.** **Gene list of DDR Targets**

| LNCaP Specific | Common | C42 Specific |
|---|---|---|
| CHTF18 | PARP1 | ETAA1 |
| TDP2 | AEN | RNF212 |
| MPLKIP | KMT2D | GTF2B |
| UTS2R | | BMP15 |
| H2AFX | | MBD4 |
| CDX4 | | PIF1 |
| PRPF19 | | FAAP24 |
| ALDH1A1 | | MDC1 |
| XRCC3 | | TTK |
| TONSL | | BRCA1 |
| TBXT | | FANCC |
| TYMS | | ERCC3 |
| RD3 | | WDR43 |
| MPG | | CLK2 |
| TREX2 | | RB1 |
| ELOB | | RNASEL |
| ERCC1 | | RECQL4 |
| POLG | | ATM |
| SHPRH | | FMR1NB |
| KLK12 | | GADD45A |
| ALDH3B1 | | |
| EME2 | | |
| WRNIP1 | | |
| APLF | | |
| HES1 | | |
| RCC1 | | |
| GRB2 | | |
| MCM5 | | |
| FAM35A | | |
| FANCG | | |
| HERC2 | | |
| CDC25B | | |
| RFC2 | | |

Supplementary Figure 4. **Comparative analysis of depleted gene overlap and DDR pathway analysis between LNCaP and C4-2 cell lines**. **A-B.** Maximum Likelihood Estimation (MLE) plots showing distribution of Beta Deviation in LNCaP (A) and C4-2 (B) CRISPR screens. **C.** Bar plot showing significantly enriched Hallmark pathways among negatively selected genes (negative RRA, $p < 0.05$) in LNCaP cells. **D.** Bar plot showing significantly enriched Hallmark pathways among negatively selected genes (negative RRA, $p < 0.05$) in common genes between LNCaP and C4-2 cells. **E.** Bar plot showing significantly enriched Hallmark pathways among negatively selected genes (negative RRA, $p < 0.05$) in C4-2 cells. **F.** Individual negative-selection gene sets (RRA, $p < 0.05$) from LNCaP, C4-2, and their overlap were independently subjected to pathway enrichment analysis.

#### Supplementary Figure 5

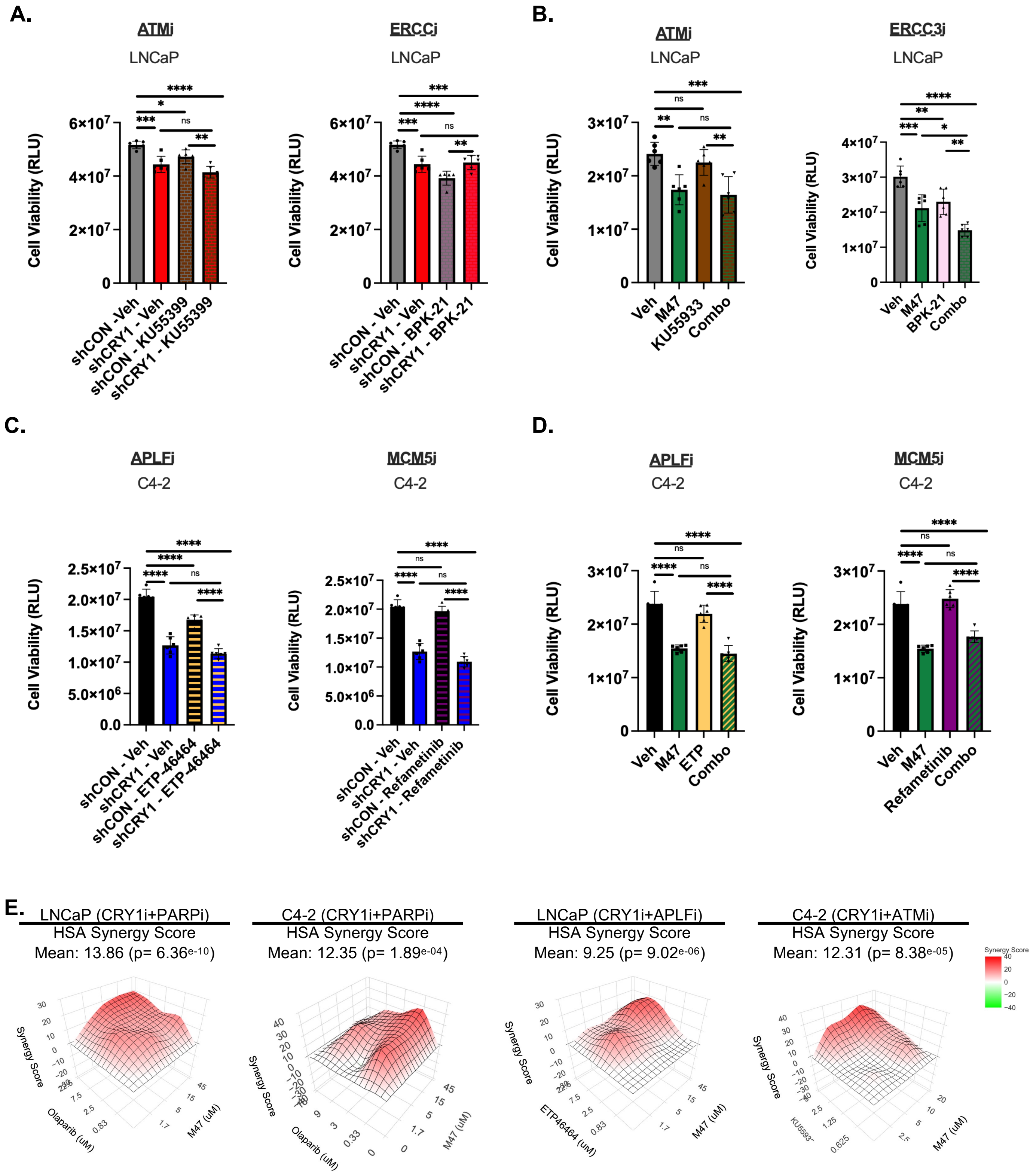

#### Supplementary Figure 5

Supplementary Figure 5. **CRY1 Inhibition Differentially Sensitizes HTS & CRPC Models to Different DDR inhibitors in a Context Dependent Manner. A.** Relative viability of LNCaP cell with CRY1 knockdown following treatment with ATM and ERCC3 inhibitors, class of DDR inhibitors unique to CRPC models, assessed by CellTiter-Glo assay. **B.** Relative viability of LNCaP cell treated with the CRY1 inhibitor M47 in combination with ATM and ERCC3 inhibitors, measured by CellTiter-Glo assay. Relative viability of LNCaP cell with CRY1 knockdown following treatment with ATM and ERCC3 inhibitors, class of DDR inhibitors specific to CRPC models, assessed by CellTiter-Glo assay. **C.** Relative viability of C4-2 cell with CRY1 knockdown following treatment with APLF and MCM5 inhibitors, class of DDR inhibitors unique to HTS models, assessed by CellTiter-Glo assay. **D.** Relative viability of C4-2 cell treated with the CRY1 inhibitor M47 in combination with APLF and MCM5 inhibitors, measured by CellTiter-Glo assay. Relative viability of C4-2 cell with CRY1 knockdown following treatment with ATM and ERCC3 inhibitors, class of DDR inhibitors specific to HTS models, assessed by CellTiter-Glo assay. All viability measurements were obtained using the CellTiter-Glo luminescence assay. Bars show mean$\pm$SEM of three biological replicates. Statistical comparisons performed using one-way ANOVA. Asterisks indicate statistical significance between groups. *$p<0.05$, **$p<0.01$, and ***$p<0.001$ ****$p<0.0001$. **E.** LNCaP and C4-2 cells were treated with the indicated Concentrations of DDR inhibitors and/or the CRY1 inhibitor, and drug interactions were evaluated using the HSA independence model. Average HSA scores approaching or exceeding 10 indicate a synergistic interaction. Most combinations demonstrated positive synergy, with some Conditions yielding scores near the synergy threshold (e.g., ~9).

### Supplementary Figure 6

A.

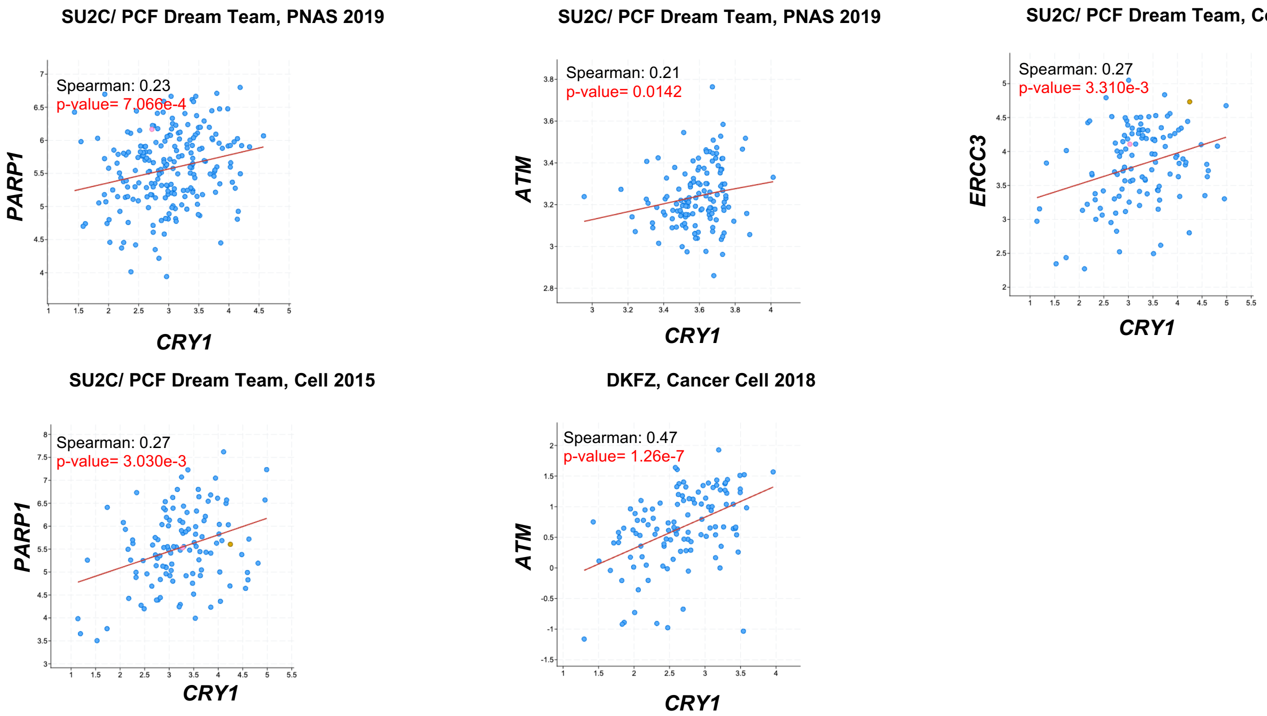


B.

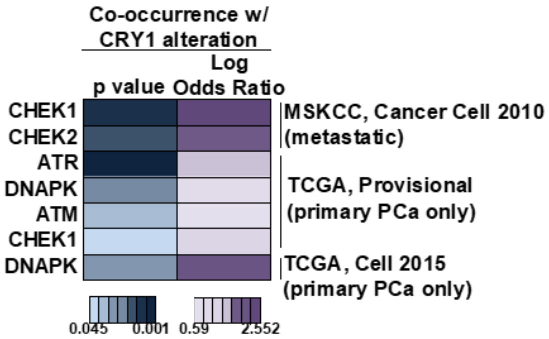


**Supplementary Figure 6**. **A.** Co-expression analysis of CRY1 with PARP1, APLF, ATM, or ERCC3 mRNA in prostate cancer (PCa) tissues. Analyses were performed using publicly available datasets, including SU2C/PCF Dream Team cohort (Cell, 2015), Su2C/ PCF Dream Team, PNAS 2019, and DKFZ (Cancer Cell, 2018). **B.** Co-occurrence of DNA damage response gene alterations with CRY1 alterations in different stages of PCa. Heatmaps show the statistical association between CRY1 alteration and alterations in CHEK1, CHEK2, ATR, DNAPK, and ATM across multiple publicly available PCa datasets. Log-transformed *p* values (left) and odds ratios (right) are shown for each gene. Analyses were performed in the MSKCC metastatic cohort (Cancer Cell 2010) and in primary PCa cohorts from TCGA (Provisional) and TCGA (Cell 2015).
